# Bridging Evolution and Design: Mapping the Diversity of LOV Photosensors

**DOI:** 10.1101/2025.09.25.678454

**Authors:** Raoul E. Herzog, Isabelle F. Harvey-Seutcheu, Philipp Janke, Wenzhao Dai, Paul M. Fischer, Peter Hamm, Philipp J. Heckmeier

## Abstract

Light-sensitive proteins allow organisms to perceive and respond to their environment, and have diversified over billions of years. Among these, Light–Oxygen–Voltage (LOV) domains are widely distributed photosensors that control diverse physiological processes. Despite their broad biological roles and increasing use in optogenetics, the functional diversity of natural LOV domains and the evolutionary constraints shaping their dynamics remain poorly resolved. A key unresolved problem is how evolution modulates the timescales and efficiencies of LOV photocycles and how this kinetic flexibility relates to biological function. Here we systematically map the photodynamics of 21 natural LOV domains – including 18 previously uncharacterized variants – and one *de novo* photosensor generated by artificial intelligence-guided protein design. We uncover an exceptional kinetic diversity spanning picoseconds to days and identify distinct functional classes within the LOV family. These patterns holistically reveal that billion years of evolutionary adaptation led to branching photocycle kinetics, matching physiological requirements. Moreover, by extending the natural catalog of LOV photosensors with a *de novo* designed LOV variant, we demonstrate how computational protein design can access new biophysical niches. This work expands the optogenetic toolkit and offers a framework to dissect and harness the evolutionary design principles of light-responsive proteins.

---

Light-sensing proteins have existed for billions of years and are found across all domains of life^1,2^. These photoreceptors are specialized proteins that enable organisms to detect and respond to light, initiating a cascade of biochemical events that enable organisms to perceive and react to their environment^3^. In recent years, humans have harnessed photoreceptor proteins for optogenetics^4,5^, a field of research that uses light to manipulate cellular functions in targeted systems^6,7^. Recent progress in optogenetics has been driven by protein engineering of photosensor domains, most notably the Light–Oxygen–Voltage (LOV) domain (Fig. 1a)^5,7–9^.

**FIG. 1.**
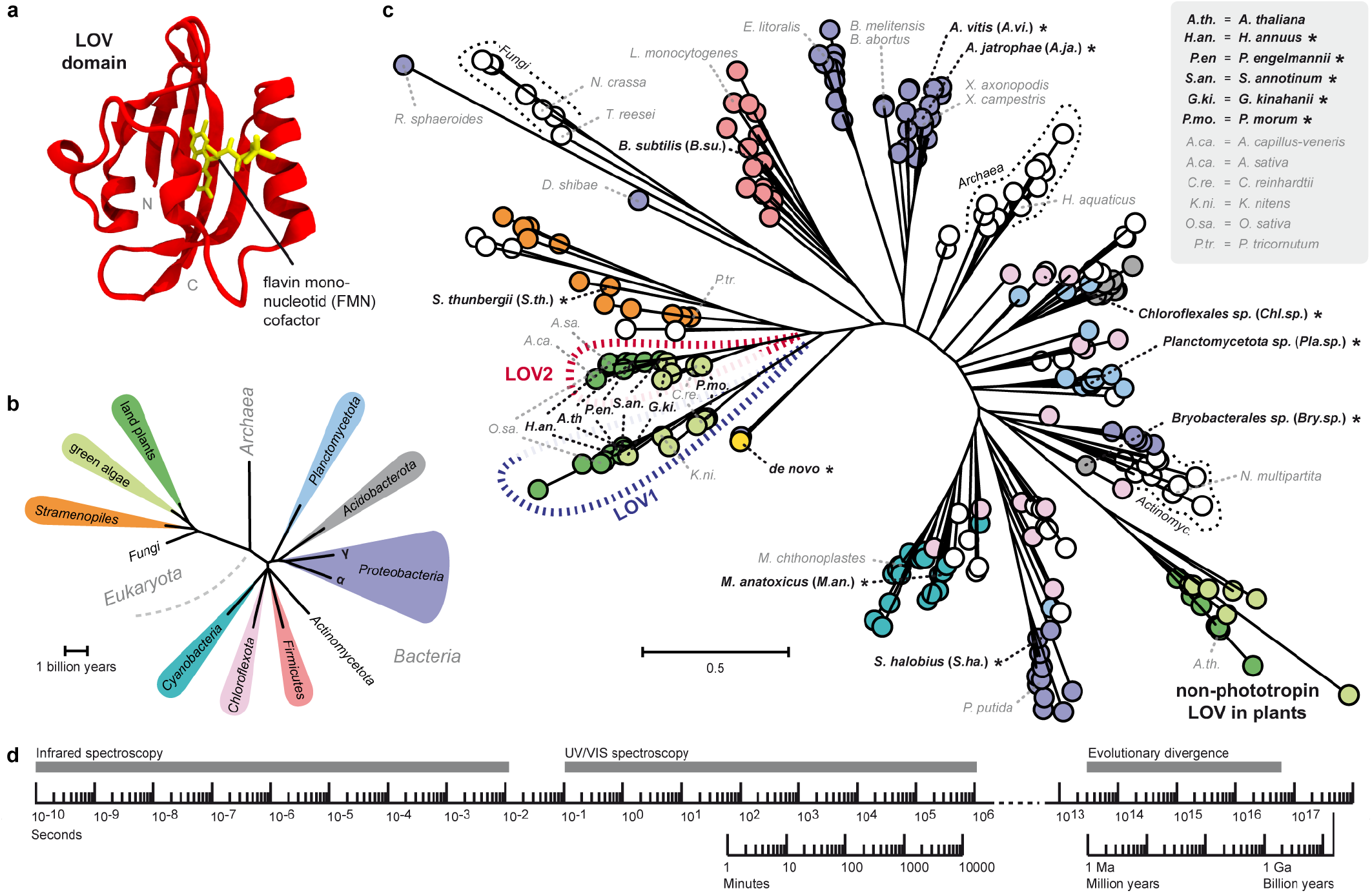
Diversity inside the LOV photosensor family. (a) Crystal structure of the LOV domain from *B. subtilis* (red) with its flavin mononucleotide cofactor (yellow) (PDB: 2PR5). (b) Distribution of LOV domains across the tree of life (TimeTree5, Feb. 2025)^20^. Exclusively LOV-containing phyla are shown; those represented in this study are color-labeled. (c) Maximumlikelihood phylogeny of LOV photosensor domains (RAxML^21^, see Methods), with highlighted variants investigated in this story (black; *: to-date uncharacterized) and already studied variants (gray). (d) Applying time-resolved spectroscopy to study the functional diversity of the billion-year old LOV protein family.

First discovered in higher plants^10^, LOV domains are blue-light receptors, known to be widespread across all branches of life, including other *Eukaryota*, numerous *Bacteria*, and even *Archaea* (Fig. 1b)^11,12^. Enabled by bacterial horizontal gene transfer, multiple LOV variants were introduced into plants, underscoring high divergence of LOV domains even within single species^11^. Reflecting this broad phylogenetic distribution, LOV domains regulate a wide variety of downstream effectors within multi-domain proteins^2^, controlling crucial processes such as phototropism, circadian rhythms, cellular stress responses, and gene regulation. Despite their abundance in the tree of life, only a limited number of LOV photosensor domains are currently deeply studied and used in optogenetic applications^13,14^.

On a molecular level, LOV domains are a subgroup of Per-ARNT-Sim domains – a primordial group of molecular sensor proteins^15^. They typically bind flavin mononucleotide (FMN) as a chromophore (Fig. 1a), which mediates structural changes upon light absorption that modulate intra- and interprotein interactions. The LOV photocycle – the core function of this photosensor domain-progresses through a series of well-defined steps: these include cofactor excitation to a singlet state, intersystem crossing to the triplet state^16,17^, formation of a covalent thioadduct between a conserved cysteine and the chromophore^14,18^ as the anchor point for further downstream signaling^19^, and the recovery to the resting, darkadapted state (Fig. 2a).

**FIG. 2.**
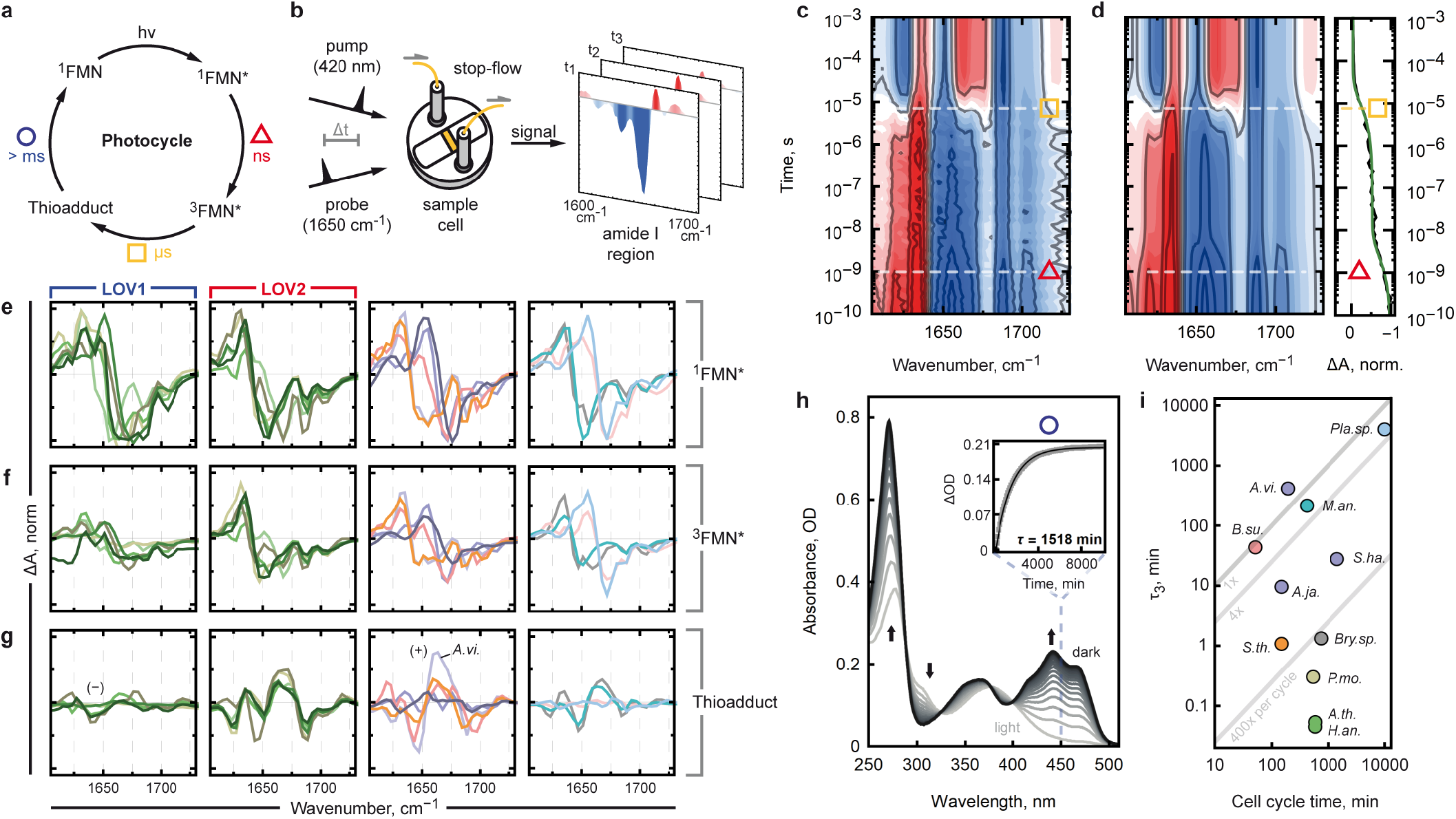
Time-resolved spectroscopy of LOV photocycle kinetics. (a) LOV photocycle upon blue-light excitation (h*?*): singlet to triplet transition via intersystem crossing (*△*), thiadduct formation (□), and dark state recovery (◯). (b) Schematic of pump–probe setup for transient infrared spectroscopy. (c) Time-resolved infrared difference spectra as a function of pump–probe delay time, exemplarily shown for *P.en*. LOV. Symbols mark time points of increased dynamic activity and the representative processes in LOV’s photocycle. (d) Global multiexponential fit of (c), obtaining exact time points of increased dynamic activity. Right panel: kinetic trace at 1650 cm^*−*1^ (raw: black; fit: green). (e–g) Evolution-associated difference spectra^40^ for ^1^FMN*, ^3^FMN*, and thioadduct. Color/species code as in Fig. 1b,c. Diverging efficiencies of thioadduct formation are marked with (+) and (*−*) and are further explained in the main text. (h) UV/VIS spectroscopy to determine dark state recovery rate *τ*_3_, exemplarily for *Chl.sp*. LOV (300 min delay between traces). Inset: Kinetic trace at 450 nm (gray) with single-exponential fit (black). (i) Recovery rate *τ*_3_ of selected LOV variants against cell cycle / replication time of their original organism (values from literature, see Tab. S3).

To dissect the molecular mechanisms underlying LOV’s function, researchers have studied mostly singular homologs from dozens of species (Fig. 1c, gray). These studies have applied time-resolved UV/VIS^16,17,22–24^, electron paramagnetic resonance^25^, and infrared spectroscopy^26–33^, X-ray scattering techniques^34–36^, and NMR spectroscopy^36,37^. However, few have taken a comprehensive, comparative perspective focused on this photosensor protein from a wide range of species. As a result, the connection between photocycle dynamics and evolutionary divergence across species remains poorly understood. What are the natural boundaries that have shaped the functional diversity of the LOV photosensors?

In this study, we aim to close this gap by providing a holistic, comparative analysis of the LOV photocycle, covering time scales from picoseconds to days, for a variety of LOV domains separated by hundreds of millions of years of evolution (Fig. 1d). To resolve early photocycle intermediates we use transient infrared spectroscopy (1) a technique highly sensitive to subtle conformational changes in both chromophore and protein, and uniquely capable of capturing ultrafast molecular events that are otherwise challenging to detect^38,39^.

By uncovering the functional variation of LOV domains, our study offers new insights into their evolutionary trajectories and provides a comprehensive overview of LOV domain diversity in the whole tree of life. Spanning both naturally occurring variants and those generated using state-of-the-art machine learning approaches, this catalog of LOV domains defines a vantage point for the modular design of new optogenetic proteins.

## SPECTROSCOPIC ANALYSIS OF 21 LOV PHOTOSENSOR DOMAINS FROM VARIOUS BRANCHES OF LIFE

In this study, we experimentally investigated 21 LOV homologs from eukaryotic and prokaryotic species, equally distributed in a phylogenetic tree (Fig. 1c, black labels). To the already existing canon of LOV domains (Fig. 1c, gray labels) that have been spectroscopically characterized, we add 18 completely unexplored LOV variants (Fig. 1c, marked with *). This selection of LOV domains is illustrated by a basic phylogenetic analysis (Fig. 1c) that aligns strongly with previous comprehensive analyses^11^. The resulting tree highlights the widespread occurrence of *Proteobacteria* variants, reinforcing their pivotal role in the horizontal gene transfer of LOV domains among all phyla of life, including early proto-eukaryotes. Furthermore, the phylogeny confirms the close evolutionary relationship between LOV1 and LOV2 domains from phototropin, a photoreceptor in green plants, while simultaneously revealing their substantial divergence from other non-phototropin LOV domains^41^ in plants. Despite the broad evolutionary separation of the 21 LOV domains in our selection, they exhibit conserved sequence motifs, particularly in residues forming the cofactor-binding pocket (Fig. S1). This conservation extends to sequence and structure similarity, as exemplified in the comparative analysis centering the *A. thaliana* variant (Fig. S2).

In subsequent experiments, we characterized the biophysical diversity among the selected 21 LOV variants, all with confirmed optical activity (Figs. S3a, S4). LOV photosensor domains undergo a photocycle, the central function of these protein domains, comprising four distinct, sequential states populated with rates in diverging time scales (Fig. 2a). We resolved three key processes: (△) the decay of the flavin singlet partially by singlet-to-triplet transition via intersystem crossing (ISC, nanosecond scale), (◻) triplet decay and concurrent thioadduct formation (microsecond scale), and (◯) dark-state recovery via covalent bond cleavage (Fig. 2a). To elucidate these key processes, we triggered the initiation of the photocycle with a 420 nm ultrashort UV/VIS laser pulse and recorded infrared difference spectra at varying delays, using time-resolved infrared spectroscopy^42^ (Fig. 2b). Focusing on the amide I region (1600 cm^−1^1700 cm^−1^, Fig. S3b), we captured spectral changes spanning seven orders of magnitude, from 100 ps (10^−10^ s) to 1 ms (10^−3^ s) (Fig. 2c, shown for *P. engelmannii*). Contour plots (Fig. S5) revealed negative (blue) and positive (red) absorption changes indicative of structural transitions^43^.

We detected two phases of kinetic activities: (△) a short-lived feature decays within nanoseconds, corresponding to the decay of the initially formed singlet ex-cited state via internal conversion, fluorescence, and intersystem crossing^16,27^. (◻) The spectrum remains unchanged for over three orders of magnitude before major features emerges at 1-20 *µ*s, marking the formation of a covalent bond between the cofactor and a cysteine residue (thioadduct)^18^. After this stage, no subsequent spectral changes were detected, indicating that the complete light-dependent structural adaption in the LOV domain is completed concurrently with adduct formation. This is supported by the agreement of the final trace with steady-state infrared difference spectra (Fig. S3b). A supplementary lifetime analysis (Fig. S6) further supported the existence of two fast processes.

In order to extract precise time constants, we applied global multiexponential fitting to a simple sequential model (Fig. 2d and Fig. S7; details in Methods). Involving three states (^1^FMN*, ^3^FMN*, thioadduct) accurately reproduced the recorded spectra, with associated time constants *τ*_1_ (decay of the singlet state) and *τ*_2_ (thioadduct formation). The obtained kinetic parameters are summarized in Tab. S1. For *τ*_1_ a range of 0.5-3.5 ns was observed, while *τ*_2_ varied between 1.2-18.8 *µ*s. These values align well with previous literature on LOV domains (*τ*_1_ =1.6-3.3 ns^28–31,44^ and *τ*_2_ =4.2-15.3 *µ*s^28–31,44^).

To understand early events in the LOV photocycle, we analyzed evolution-associated difference spectra^40^ representing singlet (^1^FMN*) and triplet (^3^FMN*) excited states (Fig. 2e,f). Their similar spectral features allowed us to estimate the quantum yield of intersystem crossing (Φ_ISC_) from relative signal amplitudes^27^ (Methods, Eq. S2). The estimated values for Φ_ISC_ (Tab. S1) show good to excellent agreement with literature values for *B. subtilis* LOV^45,46^ and plant LOV domains^16,17,23,27,47^. Across LOV variants, we observed a strong correlation between Φ_ISC_ and the singlet-state lifetime *τ*_1_ (Fig. S8). The lifetime *τ*_1_ arises from the competing rates of intersystem crossing, internal conversion, and fluorescence decay. As fluorescence rates are largely determined by the FMN chromophore and are insensitive to the protein environment^48^, this correlation implies that variations in Φ_ISC_ are driven primarily by differences in internal conversion. Thus, internal conversion acts as a gating mechanism within the photosensor’s photocycle: slower rates prolong the singlet state and increase the probability of productive intersystem crossing. Analogously, we assessed the efficiency of thioadduct formation (Ψ_eff_, Eq. S3, Tab. S1) from adduct-state spectra (Fig. 2g), which varied widely across LOV domains – ranging from inefficient formation in LOV1 (mean Ψ_eff_ = 0.14, Fig.2g −) to high efficiency in *A. vitis* (Ψ_eff_ = 0.82, Fig.2g +).

To resolve the very late process of the photocycle, the recovery of the dark adapted state (◯), we used steady-state UV/VIS spectroscopy (Fig. 2h, exemplified for *Chloroflexales sp*.; Fig. S9). We irradiated the proteins with blue light and monitored 450 nm absorbance recovery (◯) in the dark. Striking differences emerged: the recovery rate *τ*_3_ spanned six orders of magnitude (Tab. S1). From a biophysical perspective, the vast diversity in recovery times is, in fact, not particularly surprising: Modulating the activation barrier for adduct breakage by merely 5–10 kcal/mol is sufficient to span the observed range of lifetimes. Such energetic variations can be achieved through alterations in electrostatic environments or H-bonding networks within the chromophore binding pocket – mechanisms well within the scope of natural evolutionary processes, which are driven by selective pressure and functional demand. This demand becomes evident when comparing recovery rates to generation times of corresponding organisms (Fig. 2i): some LOV variants recover fewer than ten times per cell cycle, potentially regulating rare events in cell development, such as sporulation and cell division. An extreme case is *A. vitis* LOV, which could be theoretically excited only once per cell cycle (195 min). In contrast, LOV2 from a sunflower, *H. annuus*, could recover over 13,000 times per cycle, likely facilitating rapid phototropism.

## NATURE PROVIDES AN EXTENSIVE CATALOG OF DIVERSE LOV PHOTOSENSOR DOMAINS

With full photocycle kinetics available for 21 LOV domains, we compare spectroscopically derived dynamic parameters to classify LOV domains, capturing functional niches shaped by evolution (Fig. 3). Each photosensor variant represents a unique combination of the investigated photophysical parameters (lifetimes, quantum yield, etc.) and can be placed on a “biophysical landscape” that visualizes trends and correlations between these parameters. This comparative framework provides insight into how nature has diversified LOV photosensors and defines the biophysical boundaries within which optogenetic applications can be rationally designed.

**FIG. 3.**
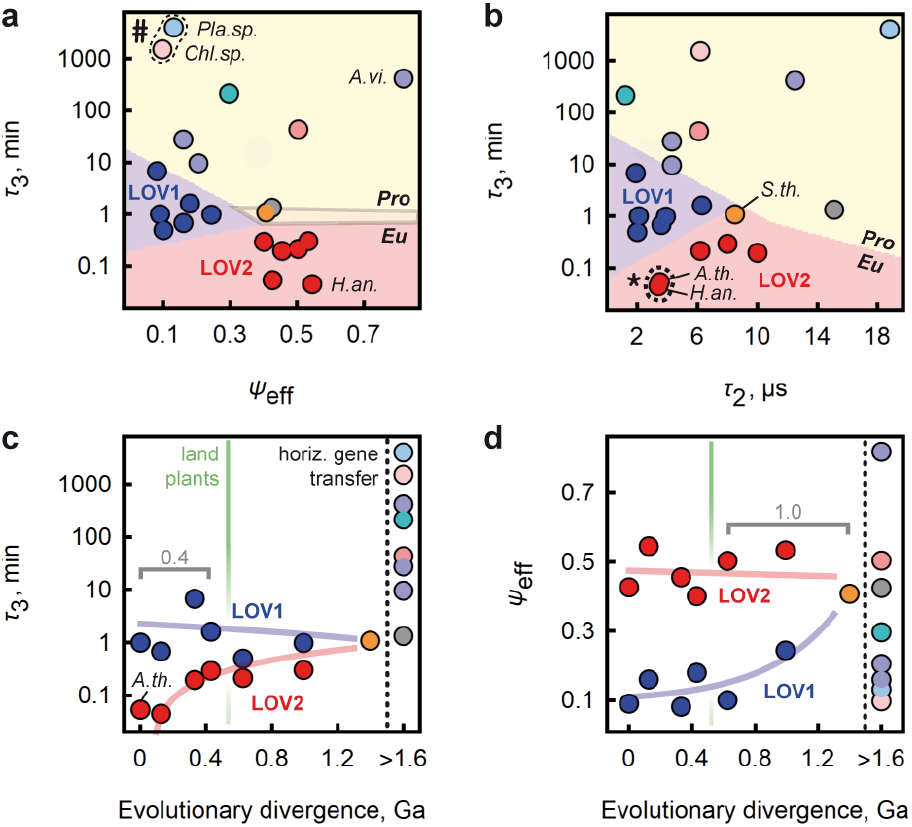
Functional classification of LOV domains. Color/species code as in Fig. 1b,c. (a) Recovery time *τ*_3_ vs. efficiency to form thioadduct Ψ_eff_. #: variants with low efficiency and ultraslow recovery. (b) Recovery time *τ*_3_ (adduct breakage) vs. adduct formation *τ*_2_. *: variants with ultrafast kinetics. For (a,b) a support vector machine analysis separates classes (see Methods), marked by shaded areas (yellow: prokaryotic LOV, blue: LOV1, red: LOV2) and by labels (Pro: prokaryotic, Eu: eukaryotic). (c) Recovery time *τ*_3_ vs. evolutionary divergence (in billion years, Ga), relative to *A.th*. LOV. (d) Adduct formation efficiency Ψ_eff_ vs. evolutionary divergence. Divergence data in (c,d) from TimeTree.org^20^ (Tab. S2), as in previous work^49^. Scales representing 0.4 / 1.0 Ga year intervals are discussed in the main text.

Comparing kinetic parameters for fast and slow photocycle processes reveals a clear separation among LOV domains where classes become apparent (Fig. 3a,b). Support vector machine analysis (see Supp. Methods) identifies boundaries that divide three classes of LOV photosensor domains: broadly dispersed prokaryotic variants with generally slower kinetics on one side, and eukaryotic domains with faster kinetics on the other side, internally dissected into two compact groups of LOV1- and LOV2-like domains.

On one of these biophysical landscapes (*τ*_3_ vs. Ψ_eff_, Fig. 3a), plant photosensors form pronouncedly distinct clusters: LOV2 domains exhibit very fast recovery and moderate to high photocycle efficiency, whereas LOV1 domains show very low efficiency and moderate to fast dark-state recovery. This divergence echoes the functional separation of LOV1 and LOV2 domains, where LOV2 retained primary photosensor function, while LOV1 likely assumed an auxiliary role, putatively as an enhancer for LOV2 photosensitivity and as a mediator for dimerization in phototropins^50,51^.

In contrast to the relatively compact plant clusters, the highly diverse prokaryotic LOV domains cover a wide range of parameters, being broadly scattered across the entire landscape. At one extreme of this plot (# in Fig. 3a), we observe both extremely low efficiency and ultraslow recovery (*Planctomyc. sp*., *Chloroflex. sp*.). On the other hand, the LOV domain from *A. vitis* efficiently reaches a very long-lived thioadduct state, making it a compelling candidate for optogenetic applications that demand both efficient adduct formation and slow adduct breakage. Another promising candidate is *H. annuus* LOV2, which combines high adduct formation efficiency with rapid thermal relaxation to the groundstate. This feature is particularly attractive for optogenetic applications that require efficient activation and precise temporal control.

The functional separation of diverse LOV is apparent but less pronounced when correlating the recovery time *τ*_3_ with the adduct formation rate *τ*_2_ (Fig. 3b). At one extreme of this functional landscape, we observe a pair of LOV2 domains from flowering plants (* in Fig. 3b) that form and break the thioadduct exceptionally fast. Other LOV2 variants show slower kinetics, and LOV1 domains are grouped separately, with significantly slower recovery times. Interestingly, the only eukaryotic non-plant LOV domain from *S. thunbergii* (Fig. 3, orange) sits directly at the intersection between the eukaryotic LOV1/LOV2 clusters and the broader prokaryotic distribution.

Independent of optogenetic perspectives, the functional divergence between LOV1 and LOV2 domains in our biophysical landscapes offers a window into the evolutionary past of plant LOV photosensors. We speculate that the functional separation between these domains happened gradually over the past 1.2 billion years, in parallel with the broader evolution of green plants. This hypothesis is supported by evolutionary trends observed across eukaryotic LOV domains, where recovery time and adduct formation efficiency correlate with evolutionary divergence (Fig. 3c,d). These relationships suggest that the two domains, co-existing in the phototropin photoreceptor, were shaped by strong selective pressures to occupy distinct functional roles. From the perspective of the flowering plant *A. thaliana*, LOV2 evolved towards increasingly faster recovery kinetics while maintaining high adduct formation efficiency. In contrast, LOV1 progressively lost efficiency in adduct formation, with stagnating recovery rates.

From the evolutionary trajectories in Fig. 3c,d, it becomes evident that the separation of LOV1 and LOV2 occurred after LOV domains were horizontally transferred into early proto-eukaryotes, but before the emergence of land plants. Based on our analysis, we estimate that this divergence spanned ≈ 1.0 billion years. The subsequent specialization of LOV2 domains in flowering plants (e.g., *A. thaliana*) toward ultrafast recovery took place later over a period of ≈ 0.4 billion years, likely reflecting an adaptation to fluctuating light environments. This biophysical tuning may have enabled finer temporal control of signaling, aligned with the ecological demands of phototropic organisms.

In conclusion, nature provides a vast catalog of LOV domains, diversified over more than a billion years into distinct functional classes. Over the course of evolution, selective pressures have driven these photosensors into highly specialized functional niches. This natural diversity presents a rich reservoir for optogenetic applications, allowing precise selection of photosensors tailored to specific needs. To go even beyond, newly-designed *de novo* proteins could extend the natural catalog of LOV domains, broadening the spectrum of potential applications.

## *DE NOVO* LOV WITH UNIQUE PROPERTIES

Building on our classification of functional clusters in LOV domains, we designed a novel *de novo* LOV photosensor and investigated its placement within the functional landscapes that were defined by key biophysical parameters in the previous chapter. Can *de novo* protein design shift the biophysical properties of the photosensor or even unlock functional niches of LOV domains that natural evolution has not yet opened?

To obtain a *de novo* LOV domain with a substantially redesigned sequence, we selected the *A. thaliana* LOV2 variant (PDB: 4EEP) as a structural template and employed the protein inverse folding model LigandMPNN for sequence redesign^52,53^ (Fig. 4a). The resulting protein adopts a predicted structure closely matching the template (backbone RMSD: 0.423Å; pLDDT: 95.9), yet shares only 51% sequence identity. Based on a calibrated relationship between sequence identity and divergence time among eukaryotic LOV domains, this places the *de novo* LOV at a hypothetical distance equivalent to 1.3 *±* 0.2 billion years from its template (Fig. 4b).

**FIG. 4.**
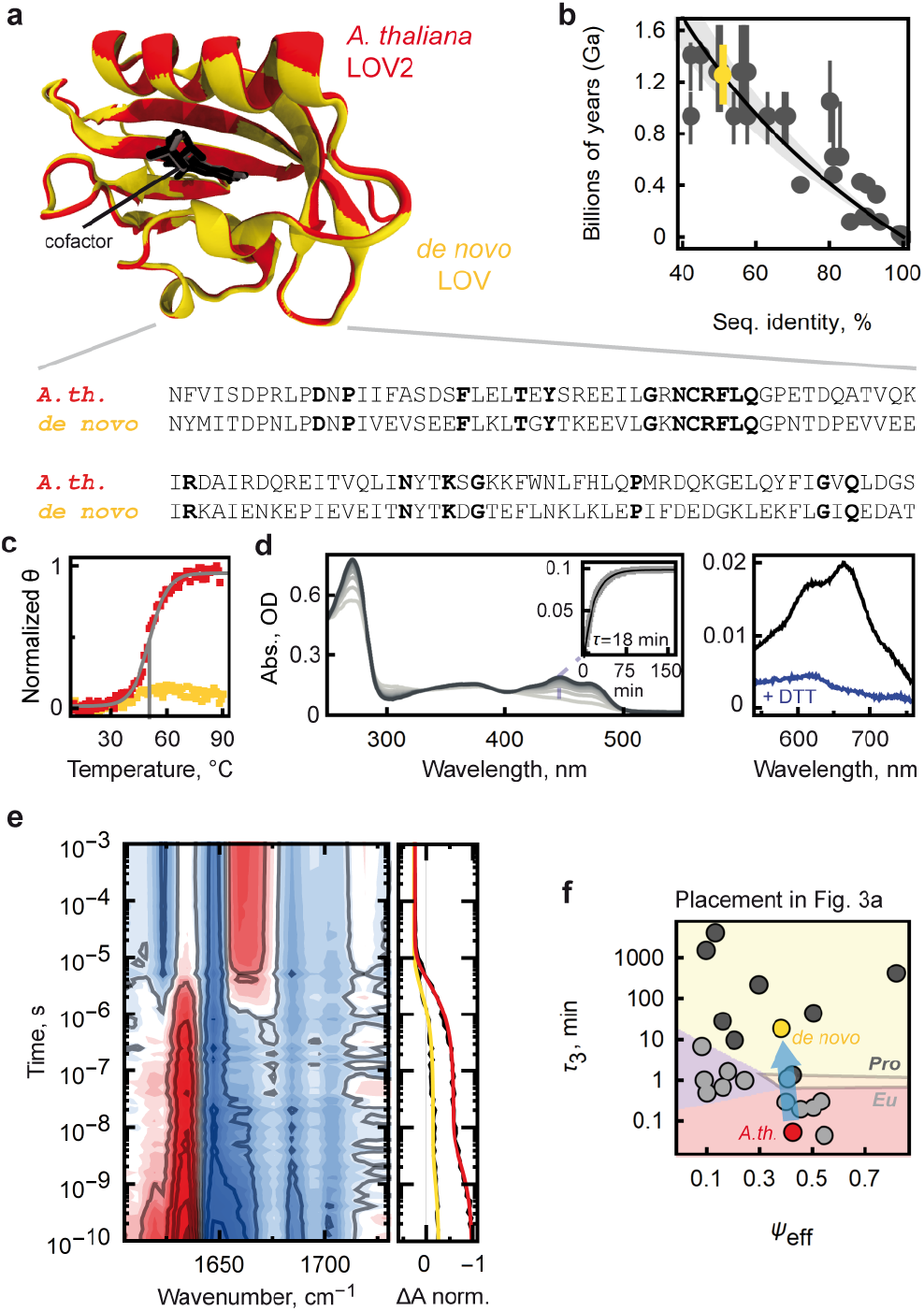
*De novo* LOV photosensor with unique biophysical properties. (a) The *de novo* LOV (yellow) was designed via a protein inverse folding model with a predefined structural output^52,53^, using *A.th*. LOV2 (red) as template. Bold: conserved residues, fixed during design. (b) Evolutionary divergence of *de novo* LOV from *A. th*. LOV2 estimated at 1.3 ± 0.2 Ga, using a logarithmic model (*y* = *−*1862.73 · log *x*) fitted to eukaryotic LOV2 sequences (gray; see Tab. S2). Shaded area: propagated uncertainty (± 0.3 Ga); local error of ±0.2 Ga at 51%. (c) Melting curves of *A.th*. LOV2 (red; T_*m*_ = 51 °C marked in gray) and *de novo* LOV (yellow); mean residue ellipticity (*θ*) at 222 nm as a function of temperature, recorded with circular dichroism spectroscopy. (d) UV/VIS spectrum of the *de novo* protein. Kinetic trace at 450 nm served to calculate recovery rate (*τ*_3_). Maxima at 600-700 nm could be diminished by adding reducing agents (dithiothreitol, DTT; Fig. S10a). (e) Time-resolved infrared spectrum of *de novo* LOV. Right panel: kinetic trace at 1668 cm^*−*1^ with global fits (yellow: *de novo*; red: *A.th*. LOV2). (f) Mapping *de novo* LOV on the functional landscape from Fig. 3a.

During protein expression, the novel protein showed extremely high yields and proved itself robust, displaying barely any aggregation or precipitation. Notably, its thermal stability far exceeds that of *A. thaliana* LOV2, remaining stably folded even at temperatures exceeding 90 °C (Fig. 4c). Performing UV/VIS spectroscopy (Fig. 4d), we observed the presence of a distinct dark/light state and a markedly slower back reaction rate, reduced by three orders of magnitude (*τ*_3_ = 18 min). A shallow absorption maximum at 600-700 nm likely originates from the formation of stable radicals at aromatic side chains interacting with the FMN cofactor^54^ (Fig. 4d). Adding reducing agent resulted in the decline of this feature, with *τ*_3_ staying unaffected in the same order of magnitude as for untreated samples (Fig. S10a). Time-resolved spectroscopy revealed that *de novo* LOV resembles its template variant (Fig. 4e), exhibiting nanoand microsecond kinetics with time constants and spectral features similar to those of other LOV2 domains (Fig. S10b,c, Tab. S1).

To contextualize our *de novo* variant within the broader functional landscape of LOV domains, we mapped its properties onto the classification framework constructed in Fig. 3a. The results, illustrated in Fig. 4f, reveal a striking functional shift (blue arrow): while originating from a member within the eukaryotic LOV2 class, the new variant resides now in the prokaryotic section, proximal to LOV domains from *Proteobacteria*. Notably, this undirected transition bypasses a cluster of points which encompasses the LOV1 domains. The extreme positioning of *de novo* LOV implies that sequential differences vitally contributed to this substantial displacement. Our approach to generate a *de novo* photosensor domain similar to *A.thaliana* LOV2, yielded a variant that preserves the ultrafast photophysical characteristics of its template while exhibiting a significantly prolonged back reaction time. With its exceptional structural stability and high expression yields, it is an excellent candidate to efficiently produce for experimental purpose, potentially in optogenetics.

Beyond that, a thrilling evolutionary insight emerges from the functional transition which we observed for the *de novo* LOV variant. Recent advances in protein design have demonstrated that artificial intelligence can generate proteins that are distantly related yet remain functional, effectively simulating hundreds of millions of years of evolution^55^. In our case, the *de novo* LOV domain is estimated to be sequentially 1.3 ± 0.2 billion years removed from its maternal template. Yet, the biophysical changes we observe do not reflect such an evolutionary leap. Unlike natural evolution, which conserves energy landscapes, structure, and functional integrity through multi-layered selection pressures, our design focused solely on fold stability. That this artificially designed *de novo* photosensor still completes a photocycle is remarkable, as it underscores the robustness of LOV domains’ core photosensor function. Moreover, evolution has selected against any designs, functional or not, prone to stable radicals near the cofactor, which we detect in the absorption spectrum of *de novo* LOV (Fig. 4d). This further illustrates how a complex interplay of selection criteria in *de facto* billion years of evolution led to immaculate molecular photosensors. While preserving LOV’s core function, evolution has produced a diverse catalog of LOV domains (Fig. 3), demonstrating its capacity to provide diversity without compromising functional integrity.

Finally, our findings emphasize the potential of rational protein design to extend the natural repertoire with engineered variants, thereby expanding the functional landscape of LOV domains and opening new roads for biotechnological applications.

## CONCLUSION

The ancient LOV protein family is ubiquitous across all major domains of life (Fig. 1), enabling organisms to perceive and respond to light, regulate circadian rhythms, and coordinate cellular processes^11^. Inspired by nature, humans have harnessed its outstanding ability to sense blue light at the molecular level, leading to innovative optogenetic applications^5^. In this study, we employed advanced spectroscopy to elucidate the functional diversity of LOV domains in nature and beyond.

By resolving the protein dynamics of more than 20 different LOV domains with the help of time-resolved spectroscopy (Fig. 2), we expanded our understanding of these molecular photosensors and their function from a comprehensive perspective. Our findings reveal an extraordinary range of kinetic processes within members of the LOV family, spanning from hundreds of picoseconds (10^−10^ s) to days (10^6^ s). Extending the already existing catalog of LOV domains by 18 to-date uncharacteristic variants, we discovered proteins with exceptional biophysical properties – such as LOV from *A. vitis* (very efficient thioadduct formation) and *H. annuus* (exceptionally fast recovery) – offering promising starting points for the development of new optogenetic strategies.

Mapping the biophysical landscape of LOV domains, we identified distinct functional niches (Fig. 3). Strikingly, we observed a clear demarcation between eukaryotic and prokaryotic variants, and between highly-specified LOV domains from plants (LOV1 and LOV2). Linking their ultrafast protein dynamics to billion-year evolutionary timescales^49^, we found that the functional divergence between LOV classes reflects evolutionary separations of approximately 0.4 billion years within the LOV2 branch and 1.0 billion years between the LOV1 and LOV2 domains.

With the help of artificial intelligence-driven inverse folding algorithms^52,53^, we engineered a novel LOV photosensor via *de novo* protein design (Fig. 4). Being hypothetically 1.3 billion years remote to its maternal template, the *de novo* LOV domain exhibits a unique combination of biophysical properties and serves as a blueprint for future approaches to create novel LOV photosensor domains in an optogenetic context. Thus, *de novo* LOV demonstrates how protein engineering can explore previously inaccessible points on the functional landscape, decoupled from evolutionary transitions that would otherwise take billions of years.

Our work underscores the importance of understanding how evolutionary processes can crucially shape protein dynamics, thereby creating new niches of protein function. By integrating biophysics with evolutionary biology, we offer a holistic perspective on molecular diversity grounded in its billion-year past and directed toward its future in anthropogenic applications, bridging evolution and design.

## ABBREVIATIONS

FMN: flavin mononucleotide
ISC: intersystem crossing
LOV: Light-Oxygen-Voltage.

## METHODS

### Bioinformatics and Phylogeny

LOV domains are abundant in the tree of life. To define the conserved core sequence of LOV photosensors, we generated a catalog of 100 LOV domains by searching proteins via standard NCBI BLAST (45-90% sequence identity). As starting points we used sequences of characterized LOV domains from *Pseudomonas putida*^56^, *Xanthomonas axonopodis*^57^, *Avena sativa* (LOV2^58^), *Aribidopsis thaliana* (LOV1^59^, LOV2^60^, Zeitlupe-type LOV^61^), *Trichoderma reesei* ^62^, *Bacillus subtilis*^63^, *Listeria monocytogenes*^64^, *Halonotius aquaticus*^65^, *Microcoleus chthonoplastes* ^66^, *Erythrobacter litoralis*^67^, *Phaeodactylum tricornutum*^68^ (all marked in Fig. 1c). The selected 100 sequences show strong sequential similarity and delivered insights into a conserved core sequence of LOV (Supporting Information, Fig. S11). For all experimentally investigated LOV domains, we retained this core interval (G26 to T127 in *B. subtilis*, PDB: 2PR5) to guarantee uniformity. Adjacent *α*-helical elements, such as J*α*, were not included in the representing core domain.

For phylogenetic analysis, the initial selection of 100 sequences was extended to 245 sequences to improve the phylogenetic resolution. The initial LOV100 selection, as well as the LOV245 selection used for phylogeny are openly available as a table and in ZENODO (the link will be provided at the proof stage). From this 245 sequences, we generated an alignment via CLUSTAL 2.1 (K-tuple size: 1, window size: 5, gap penalty: 3, number of top diagonals: 5; gap open penalty: 10, gap extension penalty: 0.05, no weight transition, hydrophilic gaps with hydrophilic residues GPSNDQERK and a BLOSUM weight matrix). With this alignment, we generated a basic maximum-likelihood phylogenetic tree, using the “build” function of ETE3 3.1.3^69^, implemented on a web-based analysis pipeline by the Kyoto University Bioinformatics Center (https://www.genome.jp/tools/ete/). The maximum-likelihood tree was computed using RAxML v8.2.11 ran with model PROT GAMMA JTT/GTR and default parameters^21^. Branch supports were calculated with 100x bootstrapping. Additionally, we included an outgroup of Per-Arnt-Sim domains from *Metazoa*, proteins far-related to LOV, which are not included in the unrooted tree, displayed in Fig. 1c. The patristic distances derived from the phylogenetic tree correlate with independently calculated BLOSUM62 similarity scores (Supporting Information, Fig. S12). The pairwise matrices for phylogenetic distances, BLOSUM62 similarity scores, and sequence identity values used in this study are available via ZENODO (accession link provided at the proof stage).

In parallel with the phylogeny based on 245 LOV sequences, we constructed a time tree (Fig. 1b) representing all phyla known to express LOV photosensors. Branches corresponding to species whose LOV domains were experimentally investigated in this study were highlighted. The time tree was not inferred from the LOV sequences themselves but was instead assembled using the *TimeTree* database (http://timetree.org)^20^, which provides median and adjusted divergence times derived from a large number of published studies. Resembling our approach in a previous study^49^, we estimated the evolutionary divergence times for selected eukaryotic species exhibiting LOV domains, with a detailed breakdown provided in Tab. S2, including times adjusted by TimeTree.org and median times with confidence intervals, exemplarily referenced to *A. thaliana*. These times reflect the most current scientific consensus (April 2025).

### Design of the *de novo* LOV

To obtain a *de novo* LOV domain with a redesigned sequence, we selected the *A. thaliana* LOV2 variant (PDB: 4EEP) with extreme fast kinetics as the structural template and employed the protein inverse folding model LigandMPNN for sequence redesign. Residues highly conserved across the majority of natural LOV domains (Fig. 4a) were fixed during the inverse folding step. Redesigned sequences with less than 55% sequence identity to the parental LOV2 were selected for structure prediction, including ligand placement, using the Chai-1 model^70^. The predicted backbone root-mean-square deviation (RMSD) and the predicted Local Distance Difference Test (pLDDT) scores were used to filter candidates for experimental validation. The final selected sequence exhibited a predicted structure highly similar to the template, with a backbone RMSD of only 0.423 Å and a high pLDDT score of 95.9.

### Protein preparation

Selecting LOV photosensor domains for the spectroscopic analysis, our criteria of choice were guided by equidistant representation from distinct branches across the tree of life. From these abundant LOV sequences, we could successfully generate 22 proteins for spectroscopic experiments, including a *de novo* variant (Tab. S3).

Examining protein function and dynamics, we heterologously expressed the LOV domains in *Escherichia coli*, using a riboflavin synthase-depleted C41 (DE3) strain (CpXribF; manX::ribM ribC::cat-ribF; provided by Tilo Mathes *et al*.^71^). Chemically-competent cells^72^ were transformed with pET-30a(+) vectors (synthesized by GenScript, NJ, USA), applying a standard heat-shock protocol. After transformation, the cells were grown in lysogeny broth medium including kanamycin and riboflavin, at 37 °C to reach an OD_600_ = 0.6-0.8. Subsequently, expression was initialized by adding 1 mM Isopropyl-*β*-D-thiogalactopyranosid. Afterwards, the expression culture was incubated at room temperature under constant shaking at 150 rpm. The expression was stopped after 20 h when the cells were harvested via centrifugation (3000 x *g*) and lysed via sonication (20 kHz, 4 x 1 min pulses). To isolate the target LOV domains from the raw cell lysate, we used an N-terminal poly-His tag and a Ni-affinity chromatography. To ensure spatial separation between the poly-His tag and the core LOV domain, we included a short GGS linker between polyHis and C-terminal target protein. The integrity of target proteins was controlled by mass spectrometry. For subsequent experiments, the samples were provided in “LOV sample buffer” (50 mM Tris-HCl, pH 8, 125 mM NaCl). The proteins were kept at -80 °C for long-term storage.

### Spectroscopy

#### UV/VIS spectroscopy

Steady-state UV/VIS absorption spectra were recorded for the selected LOV domains using a spectrophotometer (Shimadzu UV-2450, Japan). Spectra were acquired in the 200–600 nm range for both the light and dark-adapted states. In some cases, the delay between illumination (450 nm, *>*30 seconds) and measurement was long enough for partial recovery to the dark state, necessitating constant illumination even during acquisition to ensure accurate measurement of the lit state.

To capture the back reaction to the dark-adapted state (*τ*_3_), we monitored absorbance changes at 450 nm after blue light irradiation (450 nm, OXLasers, China) at room temperature (Fig. S9, Fig. 2h). The recovery was recorded under dark conditions upon illumination. For a subset of sensitive LOV domains (e.g., *M. anatoxicus, A. jatrophae*) prone to precipitation, prolonged measurements over several hours introduced spectral drift. These drifts were corrected by assuming minimal kinetic activity at isosbestic points (245 nm, 289 nm, 330 nm, and 560 nm), which are spectrally distant from 450 nm, thus minimizing interference with kinetic interpretation.

The absorbance recovery trace at 450 nm – computed as the difference between each time point and the initial value – was fitted to a single-exponential model *A* · (1 −*e*^*t/τ*,3^) where both the amplitude *A* and the time constant *τ*_3_ were treated as free parameters. To capture recovery kinetics spanning over six orders of magnitude, measurements ranged from 1.5 minutes for fast processes (with *τ*_3_ ≈ 3 s) to two-week acquisitions for the slowest recoveries (up to 4013 min or 2.8 days), allowing the signal to reach a plateau at approximately (≈ 4 *× τ*_3_).

#### Transient infrared spectroscopy

Transient infrared spectra were measured using a 100 kHz Yb-doped fiber laser/amplifier system (short-pulse Tangerine, Amplitude, France) together with an optical parametric amplifier (Twin STARZZ, Fastlite, France) with a subsequent frequency mixing stage in a lanthanum gallium silicate crystal as probe and reference pulses. The pump pulses were obtained by an electronically synchronized (Phasetech, Madison WI, USA) frequency doubled Ti:Sa oscillator (Spectra Physics, Milpitas CA, USA), to give pump pulses centered at ≈ 420 nm with a typical energy of 4 ≈ *µ*J and a temporal FWHM of ≈ 120 fs. The pump pulses were rotated by a *λ*/2 waveplate to give magic angle pump-probe polarizations. Pump and probe pulses were focused and overlapped in the sample (ca. 220 *µ*m and 120 *µ*m FWHM), while the reference was slightly offset (ca. 500 ≈ *µ*m). Probe and reference beams were transmitted through a spectrograph and detected with a 2×32 mercury cadmium telluride array detector (spectral resolution of 4 ≈ cm^−1^/pixel) with customized measurement electronics^73^.

The samples were provided in “LOV sample buffer” and were prepared under red light before measurements to ensure complete relaxation to the dark-adapted state. Guaranteeing high signal-to-noise ratios, the samples’ concentration exceeded 1 mmol/L in 1 ≈ mL. During the measurement, the sample was exchanged using a peristaltic pump combined with the stop-flow system^74^ between subsequent pump laser pulses. An additional flow path containing a back pressure regulator was connected after the pressure reservoir, in order to regulate pressure without adjusting the speed of the peristaltic pump. A flow cell with two CaF_2_ windows and a 50 *µ*m silicon spacer was used as the sample cell.

#### Fourier-transform infrared spectroscopy

Fourier-transform infrared spectra were measured in transmission with a commercial spectrometer (Bruker Corp., Billerica, MA, USA). Samples were kept in darkness for at least 24 h to ensure the accumulation of the dark-adapted state before measurement. To induce complete conversion of the sample, it was illuminated for 30 s with a 450 nm continuous wave laser with an output power of 30 mW (OXLasers, China). Transmission cells were prepared by pipetting 4 *µ*L of the protein solution on a CaF_2_ window. The window was equipped with a grooved (diameter 5 mm) to minimize capillary effects and ensure droplet centering. The cell was then sealed with a second window with a 50 *µ*m teflon spacer in between to ensure even sample thickness. We monitored amide I (C=O vibration, 1650 cm^−1^) and amide II bands (C-N stretching and N-H bending, 1550 cm^−1^), which are sensitive to conformational changes in both protein backbone and the cofactor.

#### Circular dichroism spectroscopy

We used circular dichroism spectroscopy to record melting curves, comparing the stability of *de novo* LOV and *A. thaliana* LOV2 domains. Therefore, we measured the samples (2 *µ*M in LOV sample buffer) in a quartz glass cuvette with a 10 mm path length with a commercial spectrometer (Applied Photophysics, UK). We increased the temperature from 10 °C to 90 °C, recorded spectra from 200 to 280 nm, corresponding to the *α*helical/*β*-sheet content of the protein sample, and calculated the mean residue ellipticity (*θ*) at 222 nm in deg · cm^2^ · dmol^−1^. In Fig. 4c, *θ* was normalized to its maximal change over the heating process. We determined the melting point (T_*m*_) of *A. thaliana* LOV2 from the inflection point of a sigmoid function (with vertical offset) fitted to the data.

### Data analysis

#### Global fitting of time-resolved spectroscopic data

Using time resolved infrared spectroscopy, we recorded 2D data sets *d*(*ω*_*i*_, *t*_*j*_) as a function of probe frequency *ω*_*i*_ and pump-probe delay time *t*_*j*_ (Fig. 2c, Fig. S5), spanning seven orders of magnitude from 10^−10^ to 10^−3^s. For each LOV domain, we applied global multiexponential fitting to the 2D dataset^75^, based on the assumption that the system consists of discrete states that interconvert and display time-invariant spectra.

We fitted the experimental data using multiexponential functions characterized by amplitudes *a*(*ω*_*i*_, *τ*_*k*_) and a global set of time constants *τ*_*k*_^76–78^:

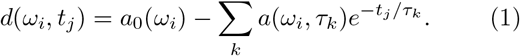

The time constants *τ*_*k*_ were treated as free fitting parameters, while ensuring that the number of exponential terms was kept to a minimum. The corresponding global fit spectra are given in Fig. 2d for *P. engelmannii* LOV2, and for all experimentally investigated LOV domains in Fig. S7. Global multiexponential fitting was applied on the unprocessed raw data of the transient infrared experiments. The only exception among all measured LOV domains was *H. annuus* LOV2, which suffered from slightly suboptimal measurement conditions. Specifically, an imperfect overlap of pump and probe beams caused cavitation resulting in an artifact (Fig. S6, marked by an *) on the nanosecond timescale^79^. In order to obtain unaffected results, the data were processed by singular value decomposition for subsequent global fitting.

The data measured with the femtosecond laser setup was globally fitted to a simple sequential model with two decaying components (^1^FMN* with *τ*_1_ and ^3^FMN* with *τ*_2_) and a non-decaying component (thioadduct)^16,17,23,27,28,32^. This model satisfactorily fitted the experimental data and provided concentration profiles of each state along with their corresponding evolution-associated difference spectra^40^, as shown in Fig. 2 e,f,g of the main article.

As reported previously^27^, spectra corresponding to ^1^FMN* and ^3^FMN* display identical features, albeit with reduced signal intensities for ^3^FMN*. Since no new signals emerge upon ^1^FMN* decay, the observed decrease in signal intensity predominantly reflects ground state recovery via internal conversion and fluorescence. In consequence, we approximated the quantum yield of intersystem crossing, Φ_ISC_, from the relative amplitudes of ^1^FMN* and ^3^FMN* evolution-associated difference spectra:

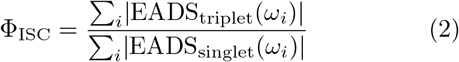

where EADS_triplet_ corresponds to evolution-associated difference spectra of ^3^FMN*, EADS_singlet_ corresponds to evolution-associated difference spectra of of ^1^FMN*, and *i* runs over the measured wavenumbers. For negative signals (bleach), the quantum yield of intersystem crossing (Φ_ISC_) can be directly estimated from the relative amplitudes of EADS_singlet_ and EADS_triplet_, without further assumptions. For positive signals, however, it must be assumed that intersystem crossing does not alter the extinction coefficients of the corresponding amide I vibrations.

While Φ_ISC_ and *τ*_1_ modulate the probability of forming the triplet state, subsequent steps within the photocycle are determined by the successful thioadduct formation. To assess this, we analyzed the spectra associated with the thioadduct state (Fig. 2g). The spectra showed substantial variation in overall intensity across all LOV domains. We quantified this observation by approximating the efficiency of adduct formation Ψ_eff_, analogously to Φ_ISC_, by

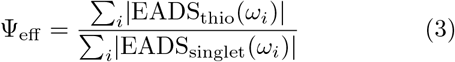

with EADS_thio_ as evolution-associated difference spectra of the thioadduct state.

Given the high predicted structural and spectral similarity among all domains (Fig. S2), the underlying structural changes and corresponding amide I difference spectra are assumed to be broadly comparable. Thus, Ψ_eff_ serves as a suitable relative metric of thioadduct formation efficiency. All acquired parameters, including the corresponding time constants *τ*_1_ and *τ*_2_, as well as Φ_ISC_ and Ψ_eff_ are displayed in Extended Data Tab. 1.

#### Lifetime analyis

Time resolved infrared spectroscopy delivered twodimensional data sets *d*(*ω*_*i*_, *t*_*j*_), defined by probe frequency *ω*_*i*_ and pump-probe delay time *t*_*j*_. In addition to a global multiexponential fitting procedure, we employed a lifetime analysis approach^75^ to independently verify the robustness of the previous global multiexponential fit, and to provide a model-free analysis of the raw data sets.

In the lifetime analysis, the time constants *τ*_*k*_ are not treated as fitting parameters but are logarithmically spaced with 10 values per decade. Only the amplitudes *a*(*ω*_*i*_, *τ*_*k*_) are optimized during the fit. To ensure stability and minimize overfitting, we applied a maximum entropy regularization scheme^75,78,80^. The resulting lifetime spectra reveal localized regions of dynamic activity, with red and blue features denoting positive and negative amplitudes, respectively (Fig. S6). To identify dominant timescales of increased dynamic activity, we calculated the “averaged dynamical content”^81^ using

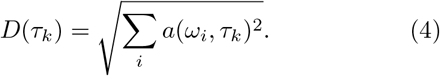

Application of the lifetime analysis to all time-resolved spectra (Fig. S5) almost exclusively reveals two dominant phases of protein response (see exception *P. morum* LOV2 in Supporting Information), consistent with the results from global multiexponential fitting.

#### Classification of LOV domains in biophysical landscapes

Mapping parameters on biophysical landscapes, acquired in spectroscopic measurements and by bioinformatics, we observed that some LOV photosensor domains share similar functional parameters, being grouped in clusters. To classify this behavior, we used a support vector machine (SVM)^82,83^ implemented using scikit-learn (v0.22.1) in Python. The dataset was split into training and test sets using an 80/20 stratified split to preserve class proportions. Prior to training, no additional feature scaling or preprocessing was applied.

A grid search with 5-fold cross-validation was performed to optimize the hyperparameters of the support vector classification (SVC) model. The search spanned the following parameter space (C: 0.1, 1, 10, 100; Kernel: ‘linear’, ‘rbf’).

The best-performing model used a linear kernel with C = 1. The optimal classifier achieved a cross-validation accuracy of 0.7619 (*τ*_3_ vs. *τ*_2_) and 0.9048 (*τ*_3_ vs. Ψ_thio_), and a test accuracy of 1.0 on the held-out test set for both datasets. The complete SVM configurations are given in the Supplementary Information.

### Usage of Large Language Models

Large Language Models (ChatGPT 4o, OpenAI, USA) were used to optimize flow of language of some sections of the main article.

## END NOTES

## Acknowledgements

We thank the lab of Alexandria Deliz-Liang for microbiological support, Thilo Mathes and Peter Hegemann, Humboldt University Berlin, Germany, as well as the Schleicher/Weber group from University of Freiburg, Germany, for the riboflavin-deficient strain (CpXribF). We thank Andreas Vitalis and Amedeo Caflisch for a fruitful discussion regarding the data analysis. We thank Serge Chesnov from Functional Genomics Center Zurich for their work on the mass spectrometry and amino-acid analysis and Roland Zehnder for the technical support. The work has been supported by the Swiss National Science Foundation (SNF) through the Sinergia grant CRSII5 213507.

## Author contributions

P.J.H. conceived the study. R.E.H., I.F.H.-S., P.J., P.M.F, and P.J.H. performed spectroscopic research. I.F.H.-S. and P.J.H. performed biochemical research. W.D. and P.J.H. performed bioinformatic research. R.E.H. and P.J.H. analyzed the experimental data. P.H. designed and built the spectroscopic setup. P.H. was in charge of funding and the research infrastructure. P.J.H. supervised the project. P.J.H. and R.E.H. prepared the figures and wrote the paper with contributions from all authors.

## Additional Information

## Supplementary Information

The online version contains supplementary material available at (link will be provided at the proof stage).

**Correspondence and requests for materials** should be addressed to Philipp J. Heckmeier.

## Peer review information

### Data Availability

The data that support the findings of this study are openly available in ZENODO (the link will be provided at the proof stage.)

