## Supporting Information for "Bridging Evolution and Design: Mapping the Diversity of LOV Photosensors"

#### CONTENTS


| | | Page |
|---|---|---|
| 1 | Supplementary Figures | 2 |
| | Figure S1 | 2 |
| | Figure S2 | 3 |
| | Figure S3 | 4 |
| | Figure S4 | 5 |
| | Figure S5 | 6 |
| | Figure S6 | 7 |
| | Figure S7 | 8 |
| | Figure S8 | 9 |
| | Figure S9 | 10 |
| | Figure S10 | 11 |
| | Figure S11 | 12 |
| | Figure S12 | 13 |
| 2 | Supplementary Tables | 14 |
| | Table S1 | 14 |
| | Table S2 | 15 |
| | Table S3 | 16 |
| 3 | Supplementary Information | 17 |
| | A. Generating LOV domains for spectroscopic analysis | 17 |
| | B. Quantum yield of intersystem crossing | 18 |
| | C. *P. morum* LOV2 shows deviating behavior | 18 |
| | D. Complete Support Vector Machine configuration | 19 |
| 4 | Supplementary Data – Predicted Structures | 20 |
| 5 | References | 21 |

### 1. SUPPLEMENTARY FIGURES

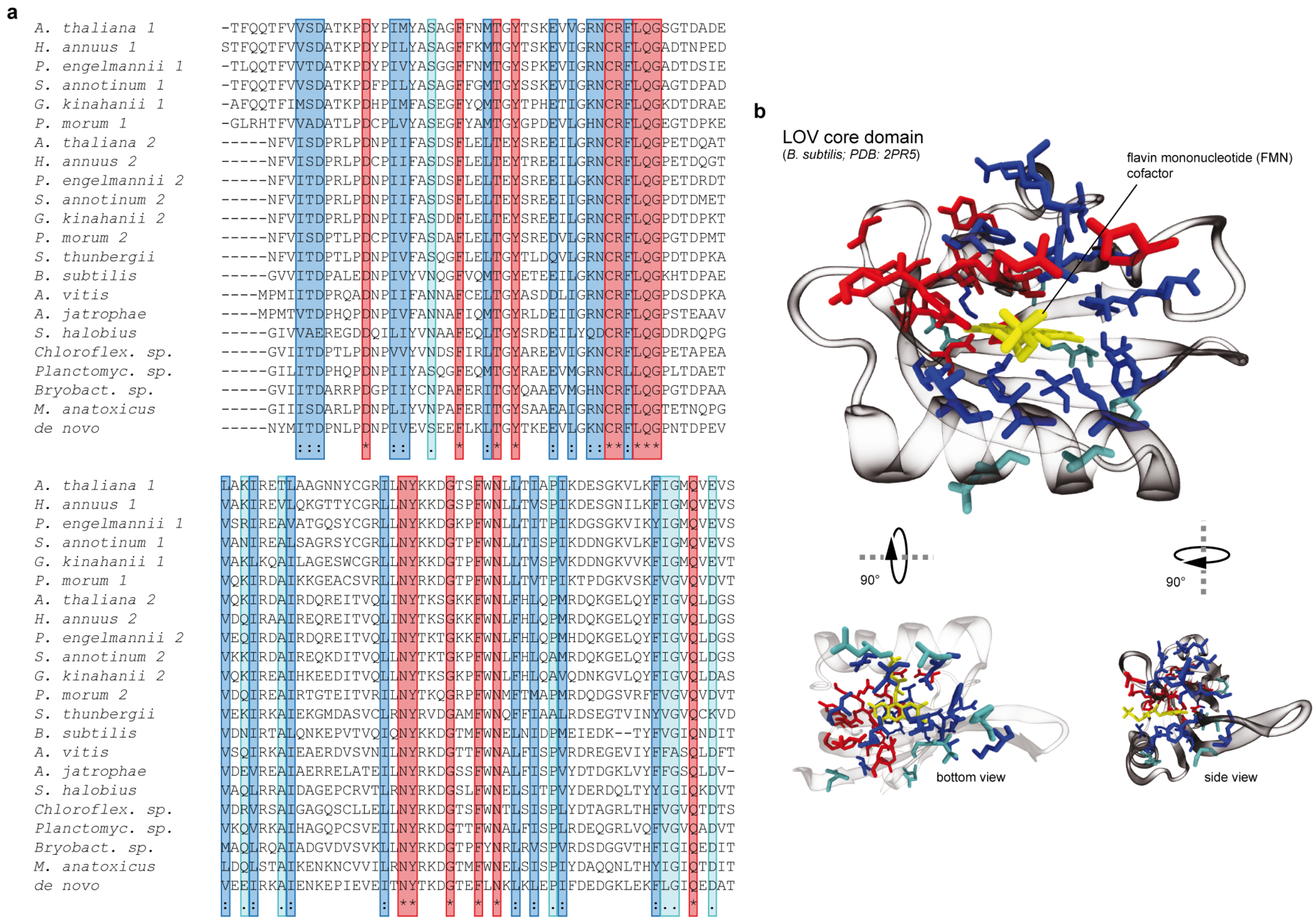


Figure S1. Selection of LOV photosensors. (a) Sequences of investigated LOV photosensor domains. "*", red: positions with a completely conserved residue. ":", blue: positions with high conservation between groups of strongly similar properties. ".", cyan: conservation between groups of weakly similar properties. (b) Location of the conserved residues in *B. subtilis* LOV.

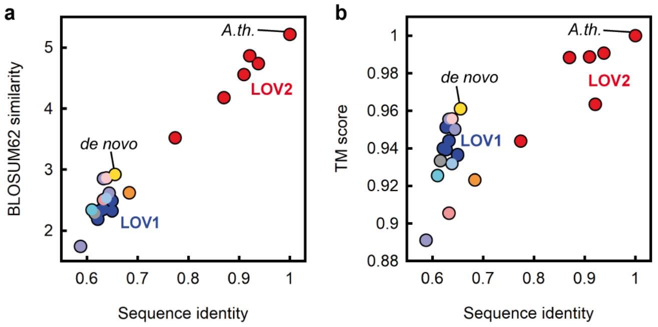


Figure S2. Comparative analysis of sequence identity (0–1 scale), (a) sequence similarity (BLOSUM62), and (b) predicted structural similarity (TM-score), all referenced to *A. thaliana* (*A.th.* LOV2). TM-scores were calculated based on structures predicted with RosettaFold[1]. Both TM-scores greater than 0.9 and sequence identities above 0.6 indicate high structural and sequence conservation among the investigated LOV domains. The color scheme as in Fig. 1b,c, with phototropin-based LOV domains are separated into LOV1 (blue) and LOV2 (red). Besides natural LOV domains, the figures also contain *de novo* LOV (yellow), generated in this study.

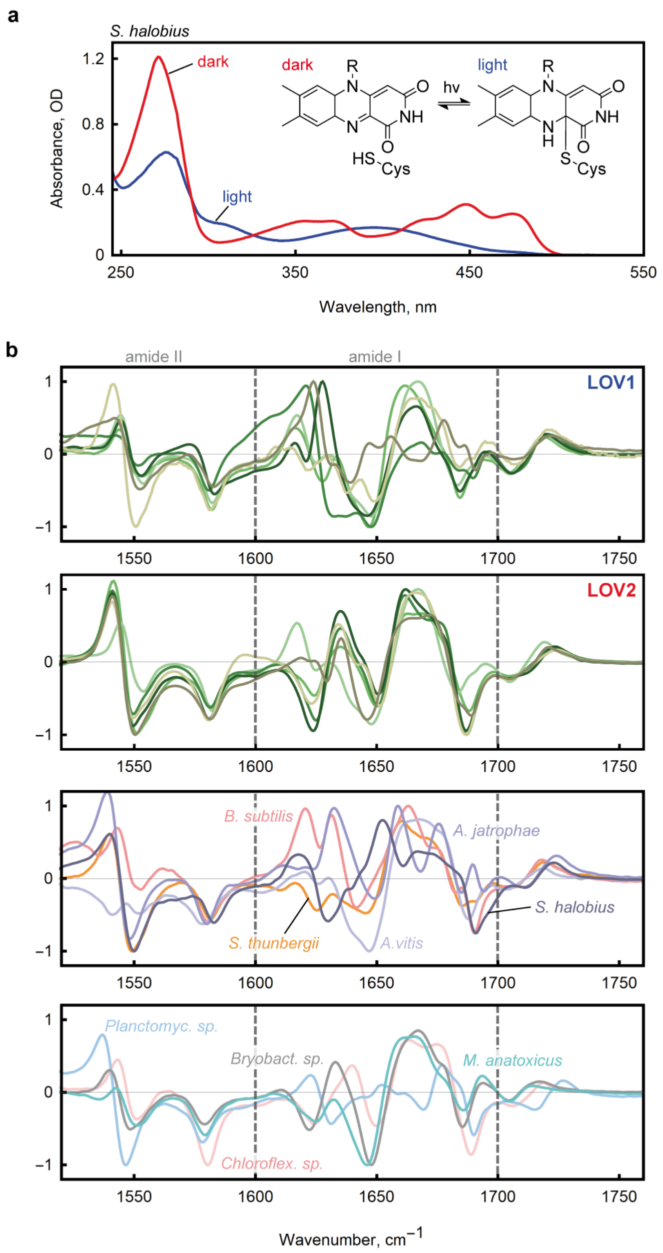


Figure S3. Basic spectroscopic analysis of selected LOV photosensor domains, in the steady-state. (a) Characteristic UV/VIS spectrum of LOV domains, here exemplarily shown for *S.halobius*. To switch to the light state, LOV was irradiated with 450 nm light. All investigated LOV domains displayed very similar spectra (Extended Data Fig. S4). (b) Fourier-transform infrared difference spectra of LOV homologs. The difference spectra were obtained by measuring fourier-transform infrared spectra in the dark and while illuminating the samples with blue light (450 nm). Color code as in Fig. **??**b,c.

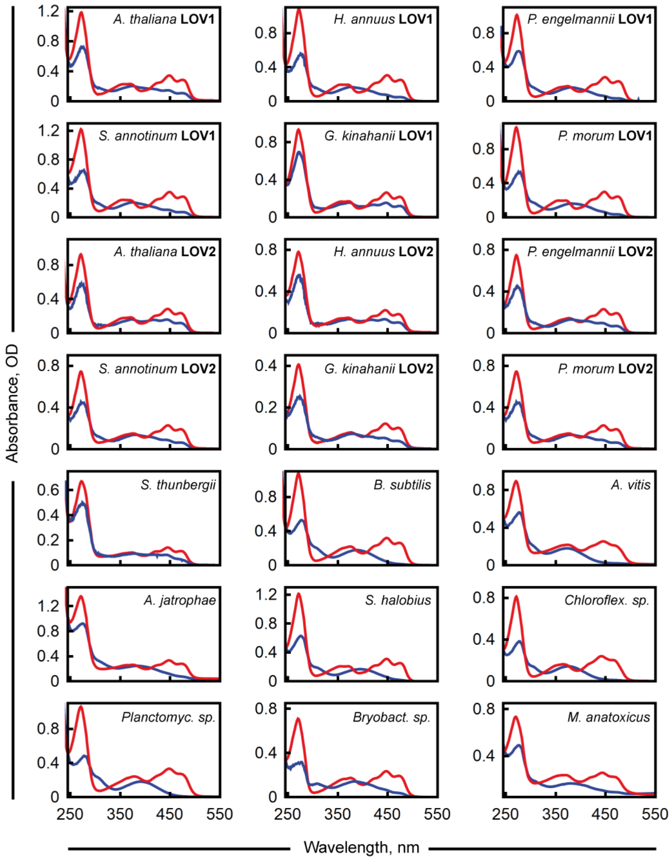


Figure S4. UV/VIS spectra of light- (blue) and dark-adapted (red) state of 21 LOV domains.

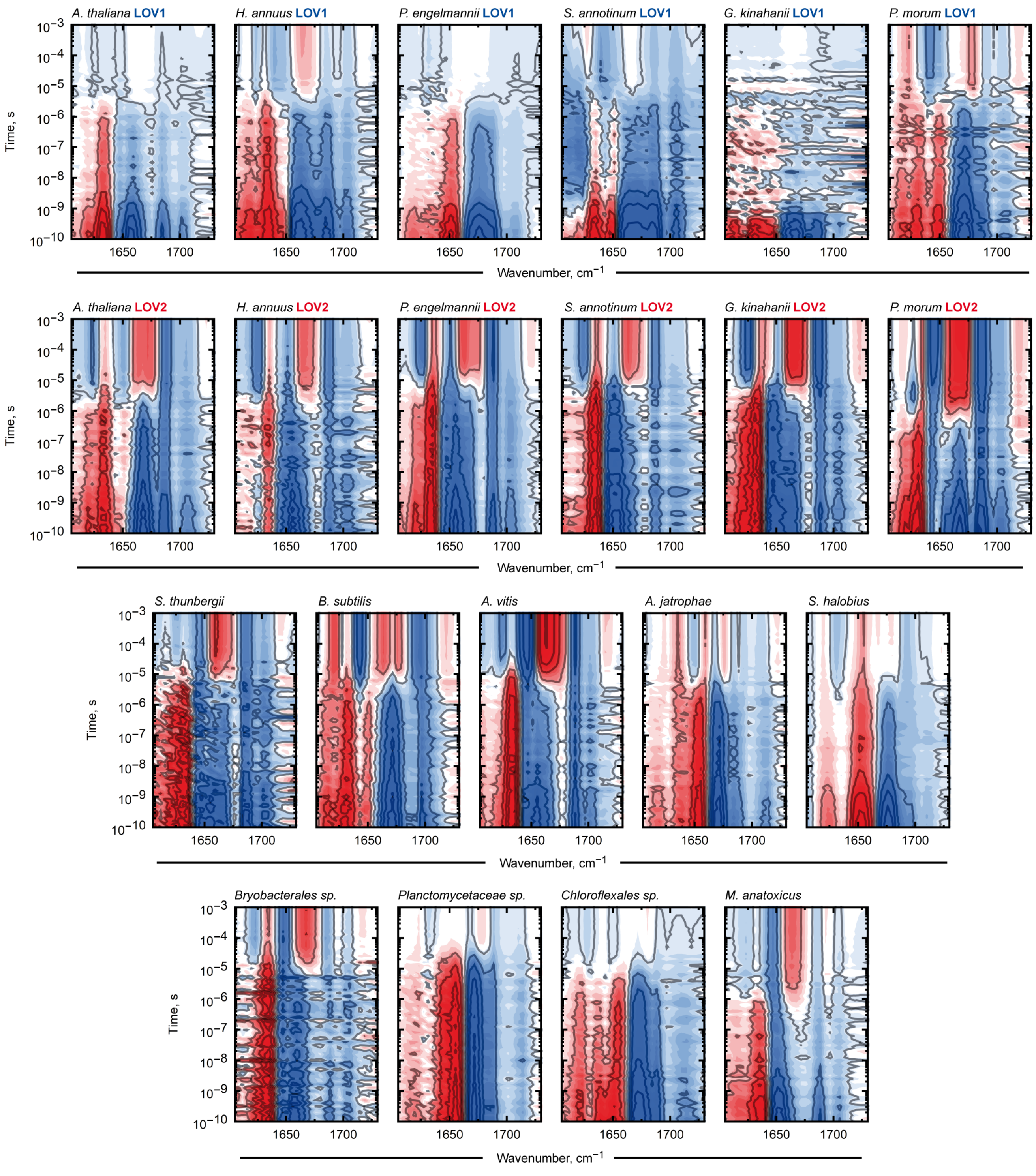


Figure S5. Raw time-resolved infrared spectra.

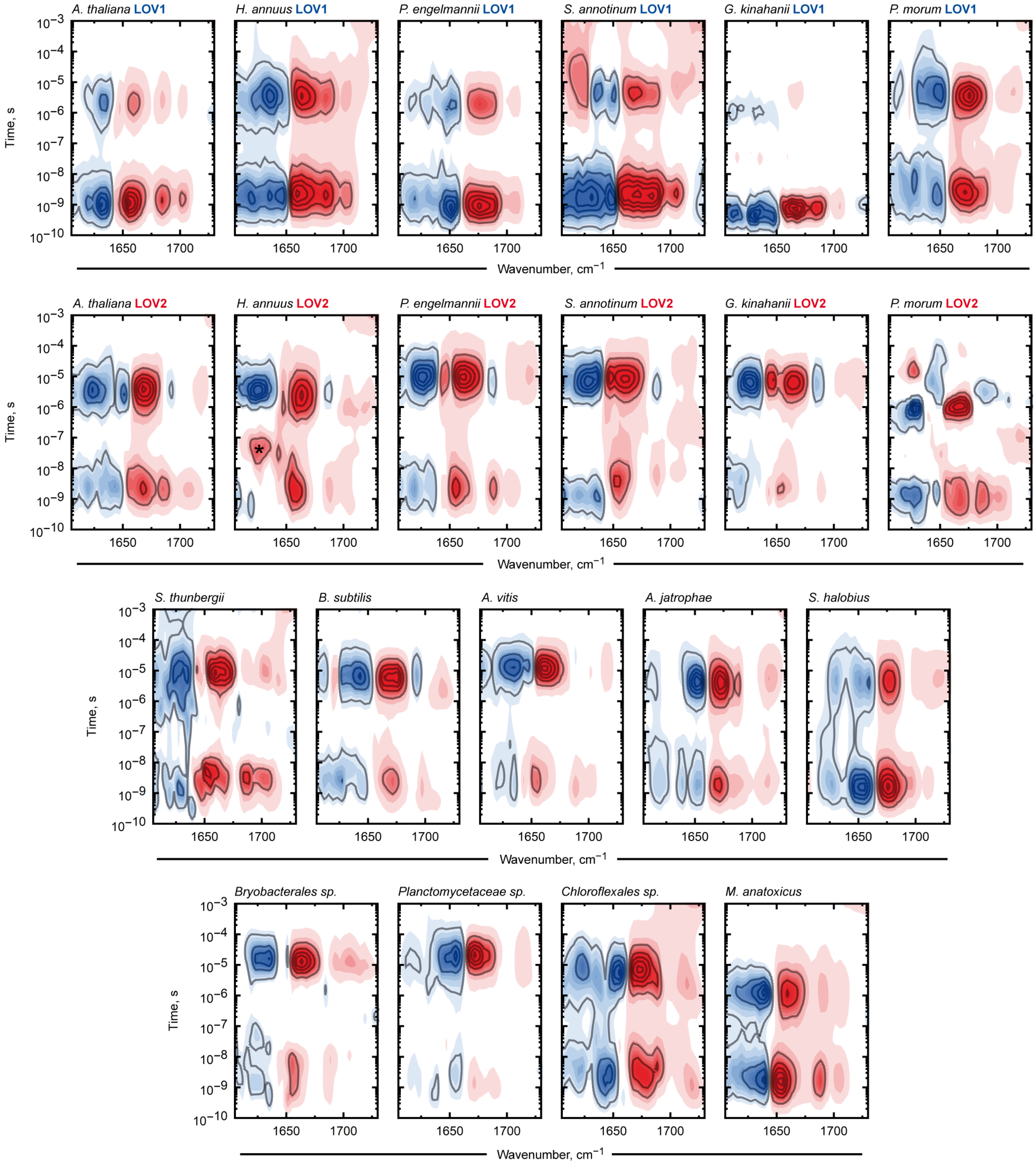


Figure S6. Lifetime analysis of raw spectra in Extended Data Fig. S5. *: artifact, see text.

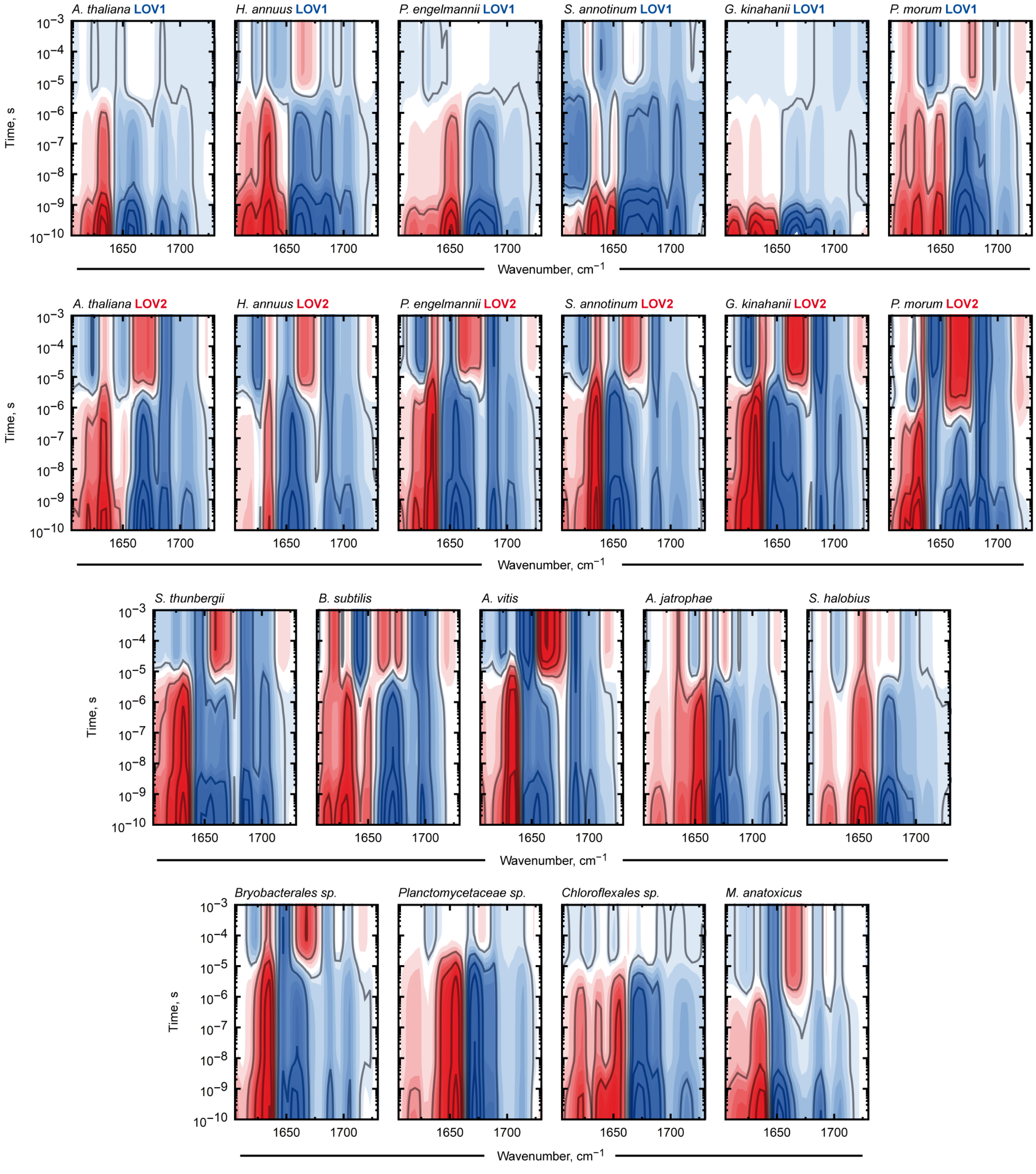


Figure S7. Global multi-exponential fitting of raw spectra in Extended Data Fig. S5.

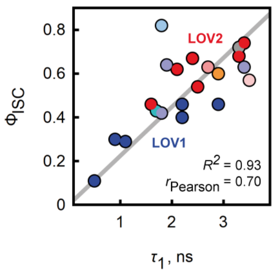


Figure S8. Intersystem crossing efficiency, $\Phi_{\mathrm{ISC}}$, vs. decay rate of the singlet excited state, $\tau_1$.

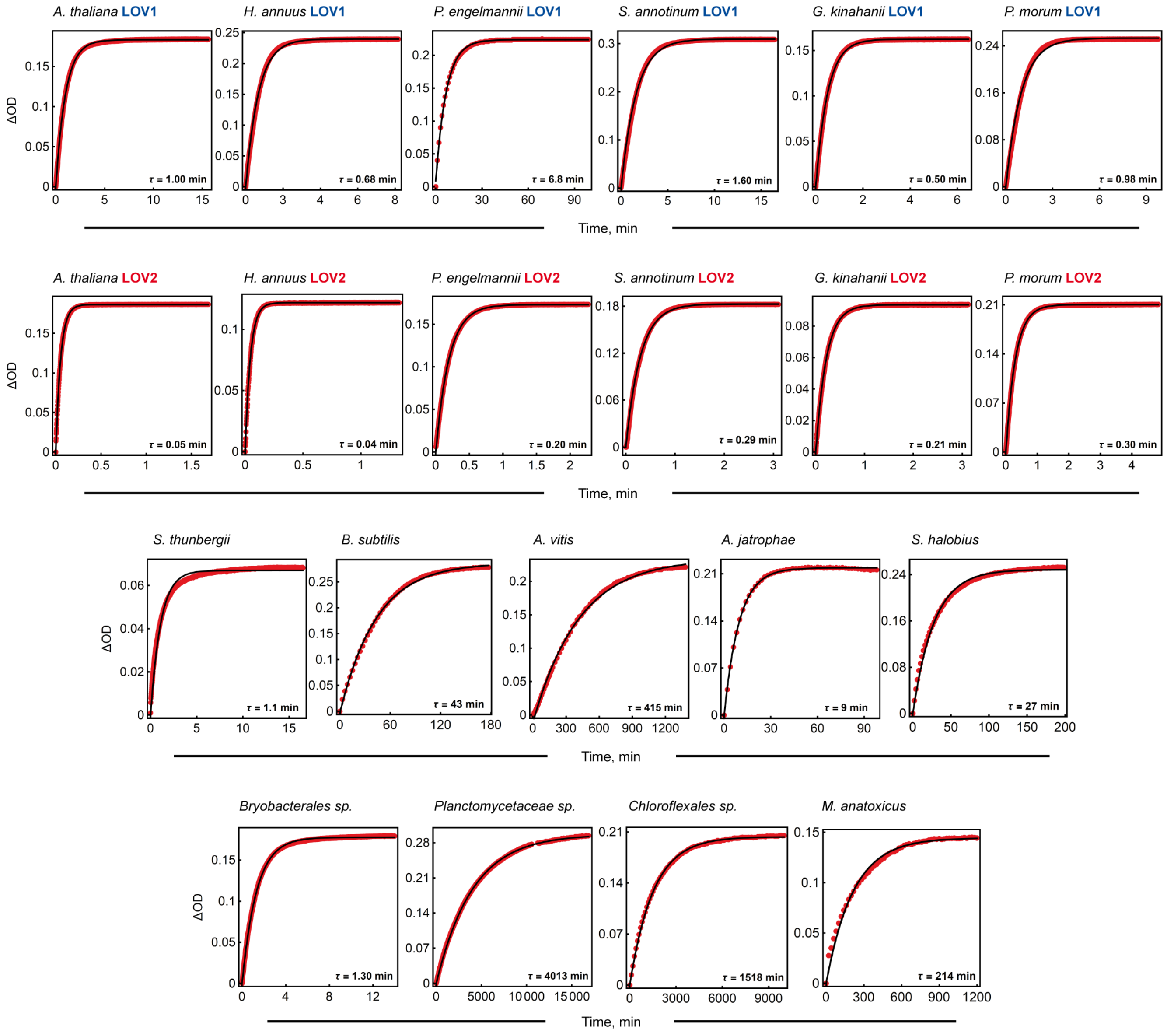


Figure S9. Dark start recovery kinetics, recorded at 450 nm (UV/VIS spectrum). Raw data (red) with single exponential fit (black).

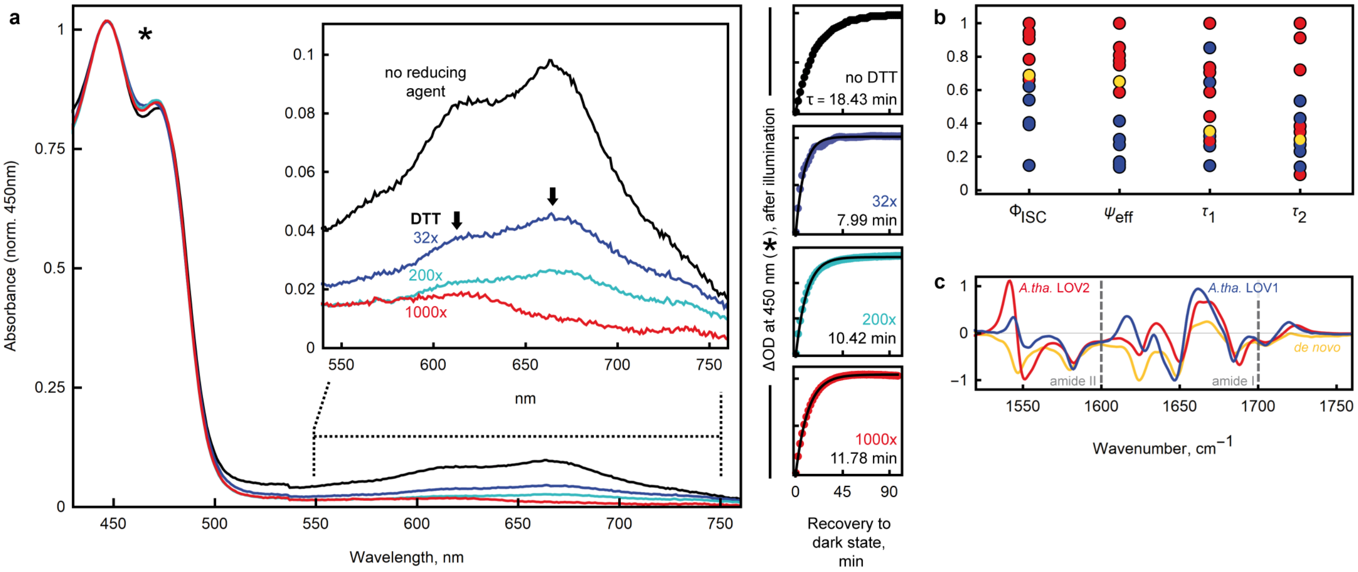


Figure S10. Spectroscopic analysis of *de novo* LOV. a) Prominent double peak at 600-700 nm that can be diminished by dithiothreitol (DTT). The addition of DTT (indicated with its excess to LOV protein; 32x, 200x, 1000x; incubation > 20 h) did not affect the order of magnitude of time constant of the dark state recovery (recorded at 450 nm, *). Interestingly, the reducing agent's effect on the double peak is heterogenous (arrows), with different behaviors for the peak at 610 nm, and the peak at 660 nm. b) Comparison of biophysical parameters between *de novo* LOV (yellow) and LOV1 (blue) and LOV2 (red), normalized to maximal value. c) Fourier-transform infrared spectrum of *de novo* LOV with *A.th.* LOV1 and LOV2 as reference.

```
                        10 amino acids
A. thaliana LOV2        NFVISDPRLPDNPIIFASDSFLELTEYSREEILGRNCRFLQGPETDQATVQK

90% consensus           shhls-....DtPllass.sF..hTGYt.t-hlGpNCRFLQs.tsp...ltp


A. thaliana LOV2        IRDAIRDQREITVQLINYTKSGKKFWNLFHLQPMR-DQKGELQYFIGVQLDGS

90% consensus           hRpulttt..hth.lhNapKsGp.hhN.h.h.Phh..tttth.hhhG.Q..h.
                                                                           4 amino acids
```

Consensus sequence legend

```
s  =  small (A, C, D, G, N, P, S, T, V)
h  =  hydrophobic (A, C, F, G, H, I, K, L, M, R, T, V, W, Y)
l  =  aliphatic (I, L, V)
-  =  negative (D, E)
t  =  turnlike (A, C, D, E, G, H, K, N, Q, R, S, T)
a  =  aromatic (F, H, W, Y)
p  =  polar (C, D, E, H, K, N, Q, R, S, T)
u  =  tiny (A, G, S)
```

Figure S11. Core domain of LOV photosensors and the 90% consensus sequence (alignment with Clustal Omega[2] with 100 sequences of the initial LOV catalog; consensus sequence visualized with MView[3]). Investigated LOV domains uniformly exhibited the displayed core domain, characterized by the interval which is spanned by (i) a virtual 10 amino acid spacer N-terminal to the highly conserved Asp, and (ii) a virtual 4 amino acid spacer C-terminal to the highly conserved Gln residue. The consensus sequence (red) was later used to construct *de novo* LOV.

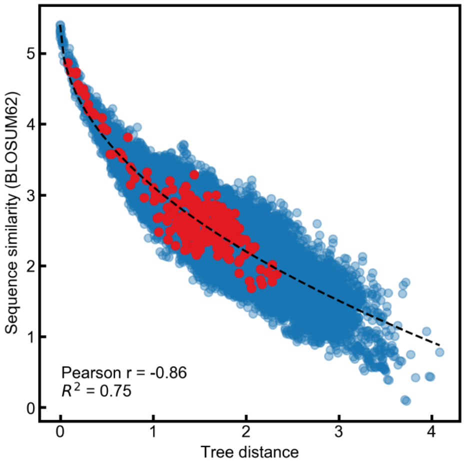


Figure S12. Sequence similarity correlates with phylogenetic distances inside the tree in Fig. 1c. The pairwise evolutionary distances (x-axis) represent the patristic distances, i.e., the sum of branch lengths connecting each protein pair in the ClustalW-generated phylogenetic tree. BLOSUM62 similarity scores (y-axis) correspond to the total log-odds substitution scores computed from pairwise sequence alignments, with higher values indicating greater sequence similarity. The plot includes all 245 sequences in the tree (blue), highlights the subset of proteins experimentally studied in this work (red), and displays a fitted stretched exponential function (dashed black; $y = 2069.7884 \cdot \exp\left(-\,(0.0000008169 \cdot x)^{0.4857}\right) - 2064.3837$).

#### 2. SUPPLEMENTARY TABLES

Table S1. Biophysical key values in the photocycle of LOV photosensor domains

| Variant | $\tau_1$, ns | $\tau_2$, $\mu$s | $\Phi_{ISC}$ | $\Psi_{eff}$ | $\tau_3$, min |
|---|---|---|---|---|---|
| *A.th.* 1 | 1.1 | 2.1 | 0.29 | 0.09 | 1.00 |
| *H.an.* 1 | 2.2 | 3.6 | 0.40 | 0.16 | 0.68 |
| *P.en.* 1 | 0.9 | 1.9 | 0.30 | 0.08 | 6.77 |
| *S.an.* 1 | 2.2 | 6.3 | 0.46 | 0.18 | 1.61 |
| *G.ki.* 1 | 0.5 | 2.0 | 0.11 | 0.10 | 0.50 |
| *P.mo.* 1 | 2.9 | 3.9 | 0.46 | 0.24 | 0.98 |
| *A.th.* 2 | 2.5 | 3.5 | 0.54 | 0.43 | 0.05 |
| *H.an.* 2 | 2.1 | 3.4 | 0.62 | 0.54 | 0.04 |
| *P.en.* 2 | 2.4 | 10.0 | 0.67 | 0.46 | 0.20 |
| *S.an.* 2 | 3.3 | 8.0 | 0.68 | 0.40 | 0.29 |
| *G.ki.* 2 | 3.4 | 6.2 | 0.74 | 0.50 | 0.21 |
| *P.mo.* 2 | 1.4 | NA* | 0.46 | 0.52 | 0.30 |
| *S.th.* | 2.9 | 8.5 | 0.60 | 0.41 | 1.09 |
| *B.su.* | 2.7 | 6.1 | 0.63 | 0.50 | 43.5 |
| *A.vi.* | 3.4 | 12.5 | 0.63 | 0.82 | 415 |
| *A.ja.* | 1.9 | 4.3 | 0.64 | 0.20 | 9.61 |
| *S.ha.* | 1.8 | 4.3 | 0.42 | 0.16 | 27.7 |
| *Bry.sp.* | 3.3 | 15.1 | 0.72 | 0.42 | 1.33 |
| *Pla.sp.* | 1.8 | 18.8 | 0.82 | 0.13 | 4013 |
| *Chl.sp.* | 3.5 | 6.2 | 0.57 | 0.10 | 1518 |
| *M.an.* | 1.7 | 1.2 | 0.43 | 0.30 | 215 |
| *de novo* | 1.2 | 2.7 | 0.51 | 0.38 | 18.4 |

We estimate an error of <10% for all parameters.

* biexponential kinetics with ambiguous assignment

Table S2. Evolutionary divergence times (in million years, Ma) of selected eukaryotic species that express LOV domains. Divergence times are based on TimeTree.org[4], ref. to *A. thaliana*.

| Species | Adjusted*, Ma | Median*, Ma |
|---|---|---|
| *Arabidopsis thaliana*# | 0 | 0 (+0.0 / -0.0) |
| *Arabis alpina* | 21 | 26 (+8.2 / -6.3) |
| *Helianthus annuus*# | 125 | 118 (+5.9 / -6.6) |
| *Mesembryanthemum crystallinum* | 125 | 118 (+5.9 / -6.6) |
| *Avena sativa* | 160 | 160 (+3.5 / -17.9) |
| *Oryza sativa japonica* | 160 | 160 (+3.5 / -17.9) |
| *Picea engelmannii*# | 330 | 330 (+6.8 / -3.6) |
| *Podocarpus coriaceus* | 330 | 330 (+6.8 / -3.6) |
| *Angiopteris evecta* | 405 | 405 (+15.7 / -8.6) |
| *Adiantum capillus-veneris* | 405 | 405 (+15.7 / -8.6) |
| *Spinulum annotinum*# | 429 | 429 (+12.7 / -19.4) |
| *Marchantia paleacea* | 484 | 480 (+17.3 / -20.0) |
| *Gonatozygon kinahani*# | 624 | 621 (+412.3 / -26.5) |
| *Cylindrocystis cushleckae* | 624 | 621 (+412.3 / -26.5) |
| *Klebsormidium nitens* | 928 | 1047 (+303.0 / -302.3) |
| *Chlamydomonas reinhardtii* | 992 | 936 (+169.7 / -207.0) |
| *Floydiella terrestris* | 992 | 936 (+169.7 / -207.0) |
| *Pandorina morum*# | 992 | 936 (+169.7 / -207.0) |
| *Pediastrum duplex* | 992 | 936 (+169.7 / -207.0) |
| *Cymbomonas sp. BC-2016* | 992 | 936 (+169.7 / -207.0) |
| *Pycnococcus provasolii* | 992 | 936 (+169.7 / -207.0) |
| *Tetraselmis chuii* | 992 | 936 (+169.7 / -207.0) |
| *Rhodochaete parvula* | 1394 | 1406 (+80.5 / -264.4) |
| *Porphyridium purpureum* | 1394 | 1406 (+80.5 / -264.4) |
| *Phaeodactylum tricornutum* | 1398 | 1278 (+348.0 / -296.3) |
| *Sargassum thunbergii*# | 1398 | 1278 (+348.0 / -296.3) |
| *Scytosiphon dotyi* | 1398 | 1278 (+348.0 / -296.3) |
| *Seminavis robusta* | 1398 | 1278 (+348.0 / -296.3) |
| *Cyclotella atomus* | 1398 | 1278 (+348.0 / -296.3) |
| *Skeletonema marinoi* | 1398 | 1278 (+348.0 / -296.3) |
| *Saccharina sculpera* | 1398 | 1278 (+348.0 / -296.3) |
| *Tribonema minus* | 1398 | 1278 (+348.0 / -296.3) |
| **Mitochondrial endosymbiosis** | 1600 | (+25.0 / -200.0)[5,6] |

### Species whose LOV domains were investigated in this study

* Adjusted divergence- and median divergence times with asymmetric errors from TimeTree.org database (accessed April 2025)

Table S3. Investigated LOV Photosensor Domains

| Photosensor | Ref. ID | Downstream Effector | Ref. | Cell cycle time, min* |
|---|---|---|---|---|
| *A. thaliana* LOV1 | 2Z6C_A | LOV - S / T Kinase | [7] | 588[8] |
| *H. annuus* LOV1 | KAJ0794400 | LOV - S / T Kinase | | 585[9] |
| *P. engelmannii* LOV1 | AML76410.1 | LOV - S / T Kinase | | |
| *S. annotinum* LOV1 | AML76953.1 | LOV - S / T Kinase . | | |
| *G. kinahanii* LOV1 | AML77647.1 | LOV - S / T Kinase | | |
| *P. morum* LOV1 | AML78527.1 | LOV - S / T Kinase | | 540[10] |
| *A. thaliana* LOV2 | 4EEP_A | S / T Kinase | [11] | 588[8] |
| *H. annuus* LOV2 | XP_021992508.1 ** | S / T Kinase | | 585[9] |
| *P. engelmannii* LOV2 | AML76410.1 | S / T Kinase | | |
| *S. annotinum* LOV2 | AML76953.1 | S / T Kinase | | |
| *G. kinahanii* LOV2 | AML77647.1 | S / T Kinase | | |
| *P. morum* LOV2 | AML78527.1 | S / T Kinase | | 540[10] |
| *S. thunbergii* LOV | AML79519.1 | bZIP domain | | 150[12] |
| *B. subtilis* LOV | WP_095843885.1 ** | STAS domain | [13] | 51.5[14] |
| *A. vitis* LOV | WP_060717380.1 ** | H Kinase (Sensory Box) | | 195[15] |
| *A. jatrophae* LOV | WP_090668721.1 | H Kinase (Hybrid sensor) | | 150[16] |
| *S. halobius* LOV | WP_109678951.1 | - | | 1440[17,18] |
| *Bryobact. sp.* LOV | MBL0157175 | - | | 750[19] |
| *Planctomyc. sp.* LOV | MBA4192485 | H Kinase (Hybrid sensor) | | >10000[20,21] |
| *Chloroflex. sp.* LOV | MBC8074770.1 | H Kinase (Hybrid sensor) | | |
| *M. anatoxicus* LOV | WP_340518864.1 ** | H Kinase | | 423[22] |
| *de novo* LOV | this study | - | | |

*Literature values for the cell cycle times of the original organism of the representative LOV

** NCBI Reference Sequence

### 3. SUPPLEMENTARY INFORMATION

#### A. Generating LOV domains for spectroscopic analysis

Out of 40 LOV photosensor domains tested for expression, 21 natural LOV variants reached expression yields above a discriminating threshold to scale up expression (target: $> 1$ $\mu$mol per sample) for time-resolved infrared spectroscopy (Fig. S13). Three additional variants reached this threshold but failed to integrate the cofactor, either *in situ* or during *in vitro* refolding experiments – most notably, putative LOV domains from *Archaea*. Six variants showed lower yields that were considered insufficient for spectroscopy; among these, one *Archaean* LOV domain also failed to integrate the cofactor. Despite these limitations, we prioritized variants from *Allorhizobium vitis*, *Microcoleus anatoxicus*, and *Sargassum thunbergii* for their importance in representing LOV domain diversity from a holistic and integrative perspective. We therefore scaled up expression of these low-yielding variants to reach the 1 $\mu$mol threshold, required for spectroscopic experiments.

LOV domains from *Xanthomonas axonopodis*, *Trichoderma reesei*, *Seiridiu cupressi*, *Rhodochaete parvula*, *Phaeodactylum tricornutum*, *Pseudomonas putida*, *Parerythrobacter jejuensis*, *Nakamurella multipartita*, *Neurospora crassa*, *Methylorubrum extorquens*, *Jackrogersella minutella*, and *Alteraurantiacibacter aquimixticola* could not be expressed under the conditions described above.

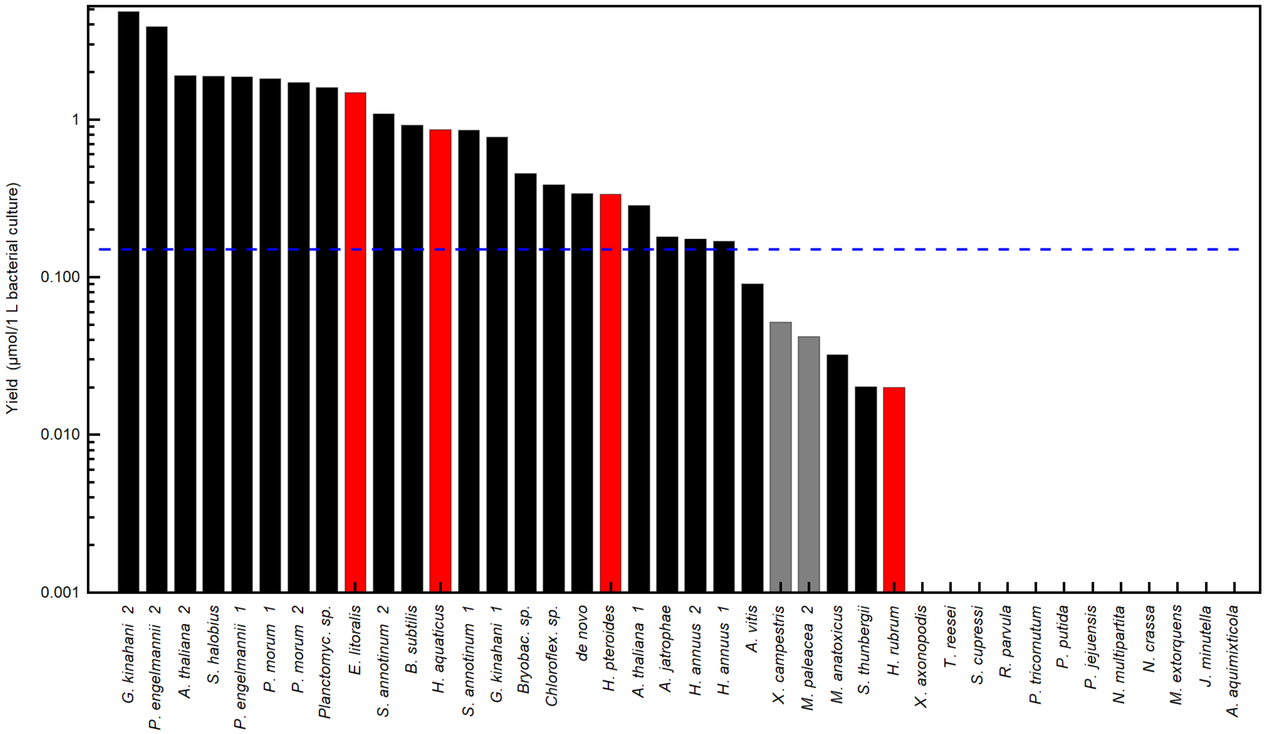


Figure S13. Expression yields of LOV domain variants, per liter of bacterial culture (in $\mu$mol), plotted on a logarithmic scale. Black bars: LOV variants that were expressed, successfully integrated the cofactor, and were included in the final spectroscopic measurements. Red bars: LOV domains that were expressed but failed to integrate the cofactor *in situ* and in *in vitro* refolding experiments. Gray bars: LOV variants that were expressed in a small amount, but whose production was not be scaled up. Dashed blue line: 0.15 $\mu$mol/L threshold used to define minimal acceptable yield.

For our spectroscopic analysis, we included land plant homologs from *Helianthus annuus*, *Picea engelmannii*, and the moss *Spinulum annotinum*, alongside previously characterized homologs from *Arabidopsis thaliana*[23]. From green algae, we included *Gonatozygon kinahanii* and *Pandorina morum.* As a non-*Plantae* eukaryotic homolog, we added the stramenopile *Sargassum thunbergii* LOV from the highly diverse SAR clade. From *Proteobacteria* we chose *Al-*

*lorhizobium vitis*, *Aureimonas jatrophae* (both *Alphaproteobacteria*), and *Spiribacter halobius* (*Gammaproteobacteria*). Other prokaryotic phyla are represented by one individual variant, i.e., *Firmicutes* by *Bacillus subtilis*[24], *Chloroflexota* by a *Chloroflexales species* from a glacier in Greenland, *Cyanobacteria* by *Microcoleus anatoxicus*, *Acidobacteriota* by a *Bryobacterales species* from activated sludge in Denmark, and *Planctomycetota* by a *Planctomycetota species* from drinking water source in Illinois (USA). Attempts to add an Archaean variant to our selection were not successful, since three expressed Archaean homologs did not incorporate their cofactor, neither *in situ* nor in subsequent *in vitro* approaches. With *H. annuus* and its pathogen *A. vitis*[25], our selection comprises two species that share the same habitat and thus the same light irradiation.

#### B. Quantum yield of intersystem crossing

We estimated the quantum yield of intersystem crossing, $\Phi_{\mathrm{ISC}}$, from the relative amplitudes of $^1$FMN* and $^3$FMN* evolution-associated difference spectra (Eq. S2). Despite this method only being an approximation, the value of $\Phi_{\mathrm{ISC}} = 0.63$ determined for *B. subtilis* LOV demonstrates excellent agreement with literature (0.62[24] for *B. subtilis* LOV and 0.62[26] for the corresponding full-length protein *B. subtilis* YtvA). Moreover, the values obtained for LOV2 (0.46-0.74) and LOV1 (0.11-0.46) domains are comparable with values in literature (0.5-0.88[27–30] for *A. sativa* LOV2 and 0.26[31] for *C. reinhardtii* LOV1). Further, we found a striking separation between LOV1 and other LOV domains: LOV1 variants consistently exhibit lower quantum yields ($\Phi_{\mathrm{ISC}} < 0.5$), indicative of reduced photoactivity, in line with previous reports[31,32].

We detected a strong correlation between the intersystem crossing quantum yield, $\Phi_{\mathrm{ISC}}$, and the decay time constant of the singlet excited state, $\tau_1$ (Fig. ED8). The efficiency of photocycle progression is governed by the competition between intersystem crossing, internal conversion, and fluorescence (i.e., radiative decay). As radiative decay is largely determined by intrinsic chromophore properties and is insensitive to the protein environment[33], differences in $\Phi_{\mathrm{ISC}}$ across LOV domains arise from variation in either intersystem crossing or internal conversion rates. The observed correlation between $\Phi_{\mathrm{ISC}}$ and $\tau_1$ suggests a model in which the internal conversion rate varies among LOV domains. A slower internal conversion rate extends the singlet-state lifetime and increases the probability of intersystem crossing. Thus, internal conversion acts as a gating mechanism at the singlet-state level – at an early stage of the photocycle, modulating how many excited molecules proceed to the productive triplet state.

#### C. *P. morum* LOV2 shows deviating behavior

Among the investigated LOV domains, one exhibited remarkably deviating kinetics: for *Pandorina morum* LOV2, the adduct formation followed complex non-mono-exponential kinetics. Analyzing the corresponding evolution-associated difference spectra (Fig. S14a), we suggest that the adduct formation completes in the early microsecond process, followed by complex additional subsequent processes that partially follow stretched exponential kinetics (Fig. S14b,c).

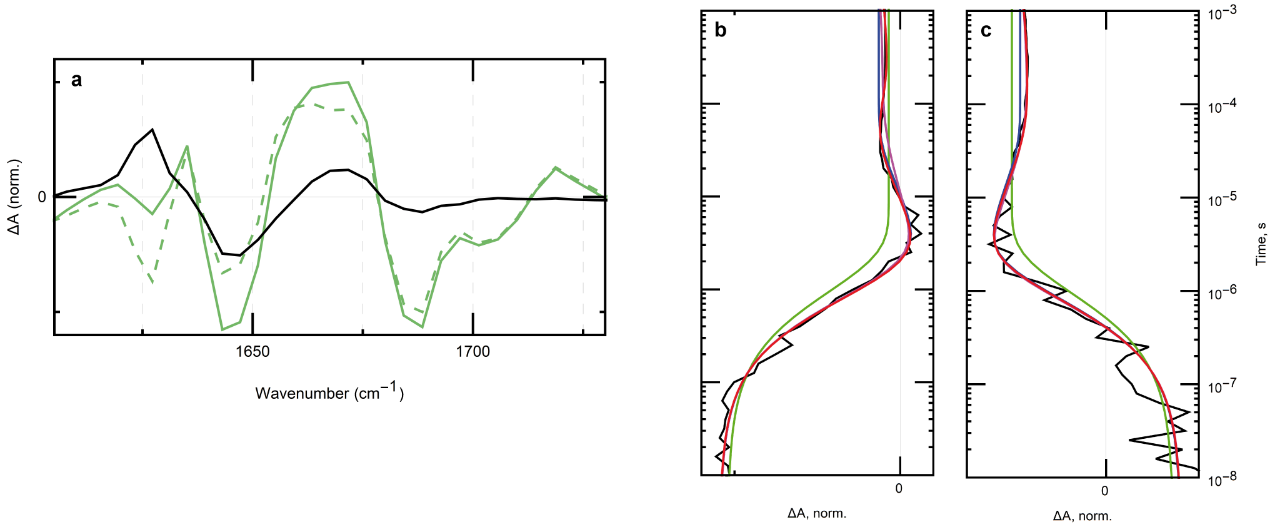


Figure S14. Deviating kinetics for *P. morum* LOV2. (a) Evolution-associated difference spectra. Green full: non-decaying state, green dashed: spectrum of the thioadduct state, black: difference spectrum (green full − green dashed). Kinetic traces at (b) 1631 cm$^{-1}$ and (c) 1655 cm$^{-1}$. Black: raw data, green: monoexponential fit, blue: biexponential fit, purple: biexponential fit with the second term stretched, red: triexponential with the second term stretched.

#### D. Complete Support Vector Machine configuration

The complete SVM configuration for ($\tau_3$ vs. $\tau_2$) are:

```
SVC (C=1, kernel='linear', gamma='scale', degree=3, coef0=0.0,
    shrinking=True, probability=False, tol=0.001, cache_size=200,
    class_weight=None, verbose=False, max_iter=-1,
    decision_function_shape='ovr', break_ties=False, random_state=None)
```

And for ($\tau_3$ vs. $\Psi_{\text{thio}}$):

```
SVC (C=100, kernel='linear', gamma='scale', degree=3, coef0=0.0,
    shrinking=True, probability=False, tol=0.001, cache_size=200,
    class_weight=None, verbose=False, max_iter=-1,
    decision_function_shape='ovr', break_ties=False, random_state=None)
```

All training and evaluation steps were performed on the same computing environment to ensure consistency.

#### 4. SUPPLEMENTARY DATA – PREDICTED STRUCTURES

The Chai-1 model[34] (status: March 5, 2025) was used to predict LOV protein structures in complex with the FMN cofactor residing in the binding pocket. The resulting files, including predicted .cif-structures, of following LOV variants are attached as .zip-files to the manuscript:

- AJatrophae.zip
- AThalianaLOV1.zip
- AThalianaLOV2.zip
- AVitis.zip
- BryobactSp.zip
- BSubtilis.zip
- ChloroflexSp.zip
- DeNovo.zip
- GKinahaniiLOV1.zip
- GKinahaniiLOV2.zip
- HAnnuusLOV1.zip
- HAnnuusLOV2.zip
- MAnatoxicus.zip
- PEngelmanniiLOV1.zip
- PEngelmanniiLOV2.zip
- PlanctomycSp.zip
- PMorumLOV1.zip
- PMorumLOV2.zip
- SAnnotinumLOV1.zip
- SAnnotinumLOV2.zip
- SHalobius.zip
- SThunbergii.zip
